# Hyper Flexible Neural Networks Rapidly Switch between Logic Operations in a Compact Four Neuron Circuit

**DOI:** 10.1101/2024.01.26.574759

**Authors:** Alexander James White, Belle Liu, Ming-Ju Hsieh, Meng-Fan Chang, Kuo-An Wu, Chung-Chuan Lo

## Abstract

Biological neural circuits at various levels exhibit rapid adaptability to diverse environmental stimuli. Such fast response times imply that adaptation cannot rely solely on synaptic plasticity, which operates on a much slower timescale. Instead, circuits must be inherently hyper-flexible and receptive to switches in functionalities without changes in network structure. This biological flexibility is a fruitful mechanism for constructing artificial reconfigurable circuits, whether they are spiking or non-spiking. In this study, we demonstrate that a 4-neuron circuit can rapidly and controllably switch between 24 unique logical functions while maintaining the same set of synaptic weights. Moreover, we show that this reconfigurability works for several different underlying neuronal architectures and strikingly can be applied to a network composed of any sigmoid-shaped activation function. We conclude with proof-of-concept applications showing that we can perform standard tasks such as a full-adder, as well as event-based conditional computing, such as detecting unexpected motion.

## Introduction

Across all organisms, nervous systems have been capable of dealing with the vast variability in their environment. Given the potentially rapid and unpredictable nature of natural stimuli, there is often no time to reconfigure a nervous system’s synaptic connections. To address this issue, environmental stimuli or contextual stimuli push the system into a different dynamical mode, that allows it to perform a different set of computations [1, 2, 3]. Likewise, there has been a recent push for reconfigurable computing, especially in the neuromorphic space, in which a circuit can be reprogrammed without changing its connectivity [4]. Biological network’s inherent rapid flexibility can be leveraged for designing reconfigurable circuits, be it neuromorphic [5, 6], organic [7, 8], or electronic [4]. Using direct inspiration from biological flexibility, we can create reconfigurable neuromorphic circuits [9].

Many studies have demonstrated this flexibility, for instance, studies have shown how small neural networks in crabs and aplysia are able to switch their function in response to different environment stimuli [10, 11, 12, 13, 14, 15, 16, 17, 18, 19, 20, 21]. Furthermore, cortical studies have shown how contextual cues can cause the networks to change activity patterns [2, 11, 16, 22, 23, 24, 25, 26]. Collectively, these phenomena of performing different behaviors based on contextual or environmental information is known as flexibility [1, 13].

Interestingly, this flexibility is important beyond just choosing the right behavior. There is evidence that flexible responses occur on a microscale [25, 27]. That is, contextual information from elsewhere in the brain will cause different responses in small collections of neurons [3, 23, 24, 25, 27, 26, 28, 29]. This flexibility extends beyond just choosing the right behavior, and can also be used in information gating [3, 16, 30, 31, 32], motor planning [10, 14, 25, 33], and influencing decision networks [13, 16, 25, 30, 31, 32]. Moreover, there is evidence that flexibility can exist on a microscale — meaning that contextual information can elicit different responses in small collections of neurons, not just in large-scaled networks [1, 3, 23, 25, 27, 29, 30].

This mechanism is noticeably absent in artificial neural networks, especially deep and convolutional neural networks (DNN and CNN). Such networks perform a single, designated task exceptionally well, but cannot generalize, as the solution hinges on a set of learned synaptic weights [34, 35, 36, 37, 38]. If the task changes, so must the weights [36, 37, 39, 40, 41]. While animals do indeed use this strategy to learn tasks, the lack of flexibility in DNNs indicates that synaptic weights alone do not explain the whole story. Many organisms are able to respond to a diverse array of situations without having to relearn a new set of synaptic weights [14, 23, 27, 42]. This is most clear in the motor cortex, where synaptic weights don’t change from task to task [10, 14, 33, 38].

Previous work on reconfigurable networks tries to endow networks with increased flexibility. Many circuits make use of memristors [4, 6, 43, 44] and homojunctions [45] to construct reconfigurable networks. These reconfigurable circuits can even be used in neuromorphic networks to implement short-time dependent plasticity and synaptic learning [4, 45, 46]. Moreover, this work has also shown that neurons are complex dynamic systems are capable of solving a plethora of tasks, especially as a network with synaptic learning [**?**, 4, 5, 6, 44, 45]. Still, none of these networks seem to make use biology’s control of the underlying dynamics of the network without (or in addition to) changing the synaptic weights [1, 14, 47, 48].

It is suspected that flexibility in the nervous system stems from its highly recurrent nature. Furthermore, these networks contain bifurcations, a qualitative change in the behavior of the system as a parameter, such as contextual input, is varied, leading to the emergence of new dynamic states [49]. Moreover, it is proposed that this is a consequence of the network being near multiple bifurcations, i.e. at a position in which state changes are easy to induce [9, 33, 50, 51, 52, 53]. Being near the cusp of state changes allows the network to rapidly switch to qualitatively different dynamics upon manipulation of certain control parameters [5, 33, 49]. However, while several studies have attempted to understand how bifurcations aid flexibility [9, 13, 54], it is still unclear how the nervous system can control networks near such complicated bifurcation structures without altering synaptic weights.

In our previous work, we identified a hyper-flexible 4-neuron circuit, named CRIREL (Coupled Recurrent Inhibitory and Recurrent Excitatory Loops). It is composed of two sets of strongly recurrently coupled neurons: a recurrently connected excitatory pair and a recurrently connected inhibitory pair, where these pairs are then weakly coupled with one another. We showed that CRIRELs are flexible because we can control an underlying double cusp bifurcation present in the network [9, 53, 48], which allows for new stable or unstable states to emerge as control parameters (in this case, synaptic weights) are varied. However, synaptic weights are not the only controllable parameters, and here we demonstrate how varying bias currents (baseline activity level, implemented as a constant background input current) and input, while keeping synaptic weights fixed, also induces bifurcations and results in flexibility. Therefore, through a detailed study of the dynamics of this microcircuit, we propose a template for how rapid flexibility can be present in neural networks without evoking slow timescale adaptation mechanisms.

In order to systematically investigate the repertoire of functionalities within CRIREL, we classify its output in terms of logic gate operations (AND, OR, etc.). These operations could be performed in terms of different types of input characteristics, and here we choose three that are often relevant in neuroscience – the difference in magnitude, timing, and phase of two input signals. In particular, timing and phase add a new type of computing, as they are event-driven, require recurrent connections, and go beyond well-trodden digital logic. To give an explicit example, during one computation, the CRIREL circuit would be comparing one aspect of two input signals, and produce a response. For example, performing an AND operation in terms of input magnitude means that the circuit will only report “on” whenever two input values have the same magnitude, and will report “off” otherwise. We found it natural to classify possible unique types of outputs given these two inputs in terms of logical truth tables (i.e. AND, OR, XOR, etc.). It is important to stress that this goes beyond just reproducing digital logic; we will demonstrate that temporal and phase processing has the capability of detecting events. Moreover, computing can be done downstream of these events. Furthermore, classifying things in terms of a logical operation has the added advantage of cleanly classifying all the different ways the network operates upon its input. We show that for all three types of input classes, we are able to generate all 8 possible nontrivial logic truth tables. Finally, we show all 24 unique functions coexist for a given synaptic weight and that changes in functionality arise solely from the change in the bias current.

## Results

### The CRIREL circuits

To examine flexibility in small neural networks, we start with a CRIREL microcircuit [Fig 1a] [9]. This microcircuit is simultaneously near two cusp bifurcations [Fig 1b] [9], and as such is capable of a simple two “bit” version of working memory [Fig 1b-c]. Here, the”bits” are loosely whether one of the inhibitory neurons is ON or is OFF. Note that two identical inputs pushed the memory circuit into a (1, 1) state, while asymmetric inputs pushed the memory circuit into a (1, 0) or (0, 1) state.

**Figure 1:**
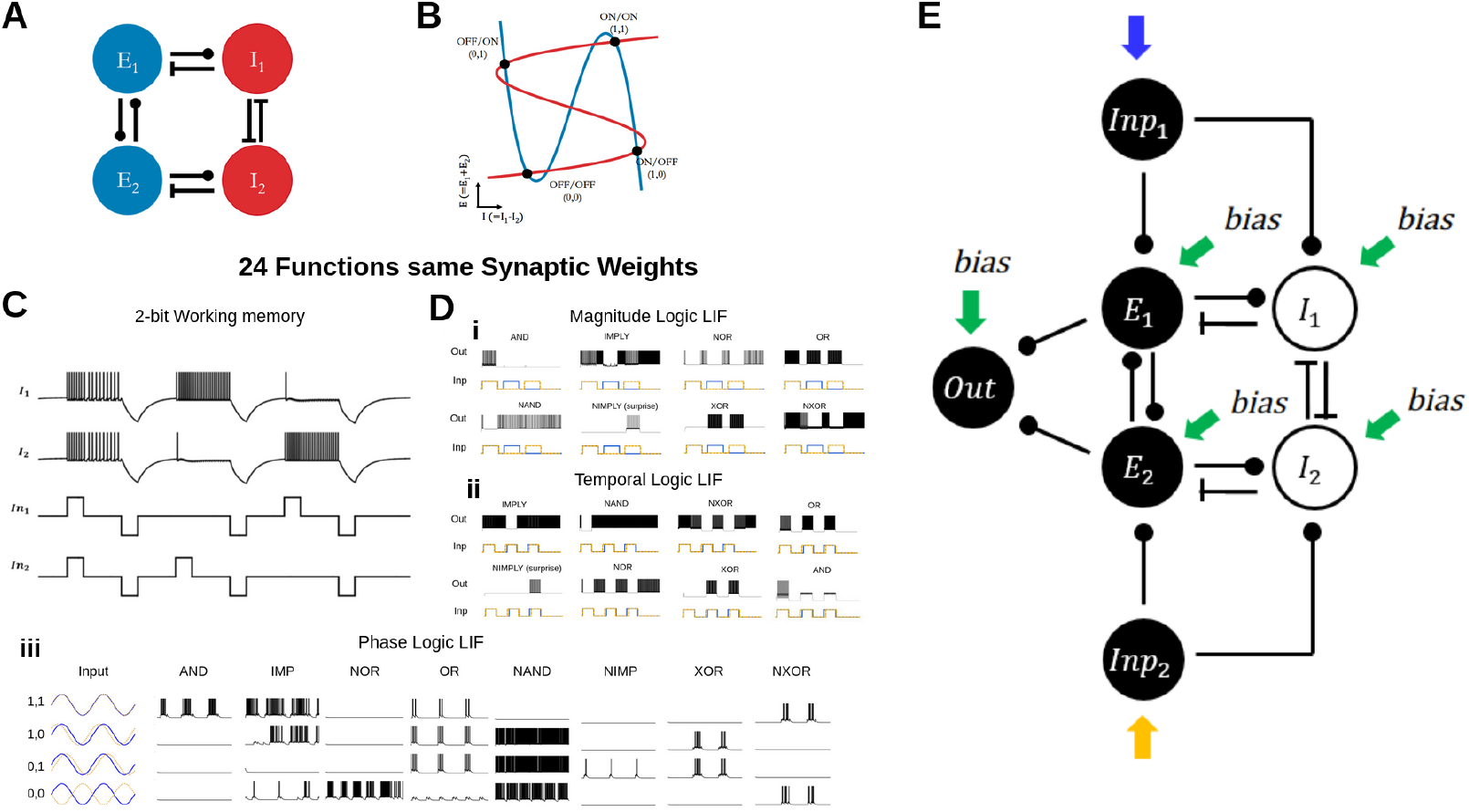
The CRIREL circuit. A) The circuit consists of two mutually coupled excitatory neurons connected to two mutually coupled inhibitory neurons. B) The 2-D reduced phase plane of the firing rate version of the network. The blue nullcline is the average excitation, and the red nullcline is the difference between inhibitory neurons. The particular setup is capable of storing 2-bit working memory C) LIF implementation of the 2-Bit Memory storage. Depending on the input into the network, the network can either store ON/ON (1, 1), ON/OFF (1, 0), OFF/ON (0, 1), or OFF/OFF (0, 0). The network can store this information until a negative reset signal. D) Twenty-four unique functions with the same synaptic weights. Only the input style and input strength change. All twenty-four distinct functions are categorized based on logic tables and input types. All Twenty-four functions share the same synaptic weight; only input type and bias current change. Di) The eight unique functions processing inputs with different magnitudes. Dii) The eight unique functions processing inputs with different timing offsets. Diii) The eight unique functions processing two periodic inputs with different phase offsets. E)The full circuit diagram to implement neuromorphic logic gates. Two input neurons are used to convert square pulses into spike trains. The output neuron is used to unambiguously map a 2-dimensional input to a 1-dimensional output. The input and output neurons are not essential for the computation of the CRIREL circuit.

In order to compute neuromorphic inputs, we add two input neurons, which convert square impulses or sinusoidal waves into spike trains [Fig 1e]. These neurons are solely for the purpose of making sure that the input into the circuit is biologically realistic. We also provide an output neuron, this is to concretely categorize the variable firing rate in the excitatory subsystem as a binary output. The output is considered one if it fires, and zero if it does not.

We compose the CRIREL circuit out of two different neuron implementation, the leaky integrate-and-fire neuron (LIF) and the Izhikevich neuron (IZH) with parameters chosen such that it is an integrator (See methods for parameters [Supplementary Table Supplementary 3 and Supplementary Table 5] [42]). Given that the LIF neuron is an integrator [49], we set the IZH neuron as an integrator rather than a resonator to provide a more apples-to-apples comparison. Moreover, this emphasizes that the flexibility arises from the network itself, not the neurons’ computational complexity.

Taking this two-bit classification as a starting point, we can classify functions in terms of binary logical operations. We take this input classification scheme, but associate the network with either an ON state or an OFF state as measured by the collective activity of the excitatory subsystem. This gives rise to 8 unique non-trivial input combinations, AND, OR, XOR, NAND, NXOR, NAND, IMP, and NIMP [Fig. 1d] (Supplementary Table 1), where the two inputs into the circuits can be classified as 1 or 0. The binary pair (1, 1) references if the inputs are “roughly” the same. Likewise, we can consider whether an out neuron is on or off as a binary output of 1 or 0. Moreover, we consider three unique types of inputs, the differences in magnitude between two square pulse inputs[Fig. 1Di], the differences in timing between two square pulse inputs[Fig. 1Dii], and the differences in the phase of two sinusoidal inputs [Fig. 1Diii]

### Magnitude Logic

The most straightforward type of input to consider is the differences in magnitude between two square pulse inputs [Fig 2a]. After being fed into the input layer, the magnitude of the square pulses is converted into a difference in firing rate, and roughly corresponds to different magnitudes of post-synaptic excitatory input. We consider the input to be (1, 1) if the magnitudes *A* are roughly equal. Likewise, if the inputs are different by a fixed amount, Δ*A* they are considered (1, 0) or (0, 1). An absence of input is considered (0, 0). It is important to note, that the magnitude *A* and difference in magnitude Δ*A* are both free parameters [Supplementary Figure 1].

**Figure 2:**
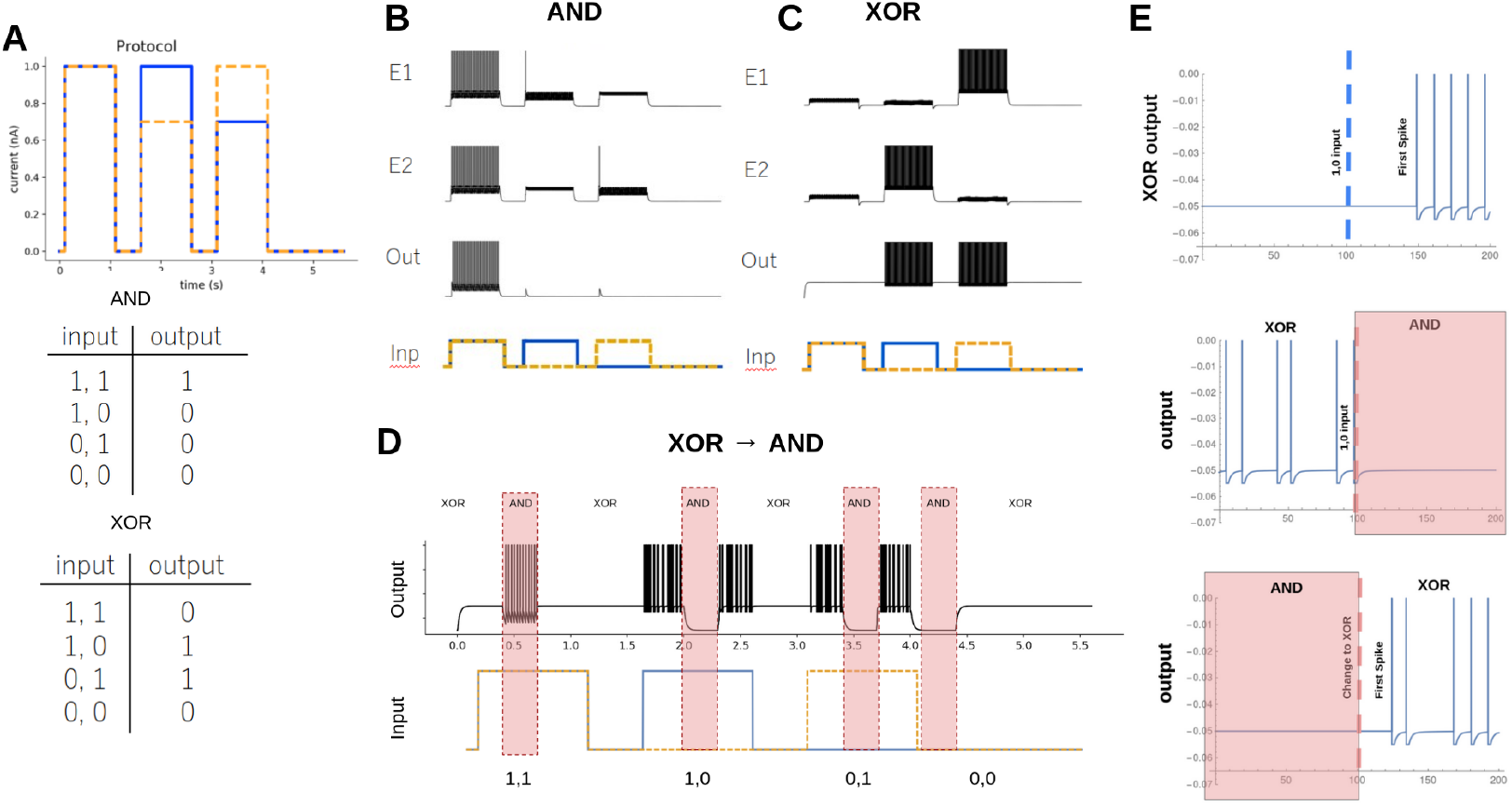
Magnitude Logic. A) Input protocol for the Magnitude Logic Gate. The difference between the magnitude of the square pulses determines the (1, 1), (1, 0), (0, 1) and (0, 0). The Truth tables for AND and XOR are shown. B) The LIF version of an AND gate. The activity of the excitatory neurons (E1 & E2) and the output neuron (Out) are shown. A cartoon representation of the input protocol (Inp) is also shown. C) Same as B but for an XOR gate. D) An example of the CRIREL being converted back and forth between XOR and AND gates on the fly during the signal input. E) The latency of switching output for the input a (0, 0) to a (1, 0) in an XOR gate is 47 *mS* or about 5 spikes. Latency of gate switching which occurs on the scales of 20 ms or about 2 spikes.

As a concrete demonstration, we begin with an AND gate. For the particular example, we choose the magnitude for 1 to be *A* = 1 *nA* for the LIF neuron. We set Δ*A* = 0.3 *nA*. When the network is presented with the input protocol, we see that the output neuron fires only when the signal is in the (1, 1) state [Fig 2b]. Simply by changing the bias currents *I*_*bias*_ (see methods), we can change the gate into an XOR gate [Fig 2c], where during the (1, 1) signal the output is silent, but fires whenever the signal is in the (1, 0) or (0, 1) signal. Next, we show how we can change the bias currents, from AND to XOR in the middle of the input protocol [Fig 2d] and the gate switches with a latency of 40 *msec* or about two spikes [Fig 2e]. A majority of the transitions are possible, but a few require global reset (strong negative input)[Supplementary Figure 2].

Next, we show that all 8 unique logic gates exist for at least some set of bias currents. We test this for LIF neurons [Fig 3a] and for Izhikevich (IZH) neurons in integrator mode [Fig 3b]. The bias currents used to produce these 8 gates can be found in the supplement [Supplementary Table 7 and Table Supplementary 8]. We provide the parameter sweep for 6 of the logic gates as a 2-dimensional sweep of excitatory subsystem bias and inhibitory subsystem bias [Fig 3c and 3d]. Because IMP and NIMP require asymmetric bias currents, they don’t fit neatly into a 2-dimensional sweep, and are not shown.

**Figure 3:**
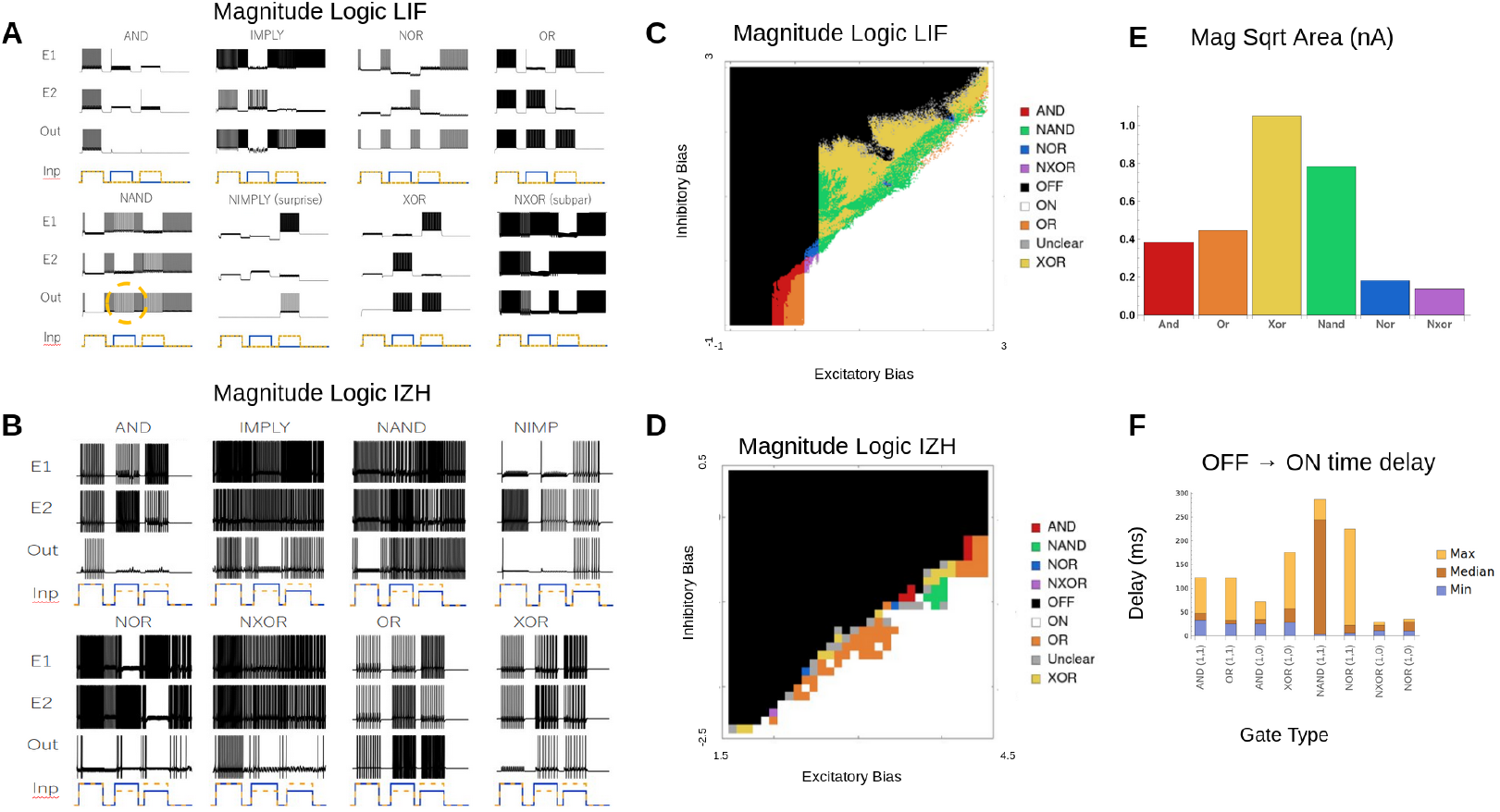
All Magnitude gates. A) All 8 Magnitude Logic gates for the LIF version of the CRIREL. Both IMP and NIMP have a mirror image that is not shown. B) All 8 Magnitude Logic gates for the IZH version of the CRIREL. Both IMP and NIMP have a mirror image that is not shown. C) A sweep through the inhibitory bias and excitatory bias for the LIF model. Each gate is labeled based on its color. D) A sweep through the inhibitory bias and excitatory bias for the IZH model. Each gate is labeled based on its color. E) The robustness of each LIF gate in terms of bias current. It is calculated as the square root of the area for each gate in Panel C. Larger values represent more robust gates. F) The average time from a silent OFF state for the output neuron to the first spike of an ON state for the output neuron. We sampled different biases from the sweep in C.

It is important to note that while each gate has a region of parameter space, there are areas where the order of inputs matters. This is because hysteresis and bistability cause the preceding input to affect the computation in the next input. As such, we mark these as unclear (gray) in the parameter space [Fig 3c and 3d].

Most of the logic gates are robust against variation in the bias currents. This robustness can be quantified as the square root of the area of each gate’s parameter region in the 2-dimensional sweep [fig 3c] of excitatory subsystem bias, and inhibitory subsystem bias [Fig 3e]. This roughly quantifies the change in bias that each gate can tolerate. Some gates, like XOR and NAND are very robust, while NXOR and NOR, require greater precision in bias current.

The gates also respond rapidly to the presentation of input. To quantify response time, we measure how long it takes for the ‘0’ (OFF) response of the output neuron to switch to a ‘1’ (ON) response. Here, we measure the time difference from when the input is presented to the network to the timing of the first spike in the output neuron. We sampled multiple different bias current configurations, some are faster than others, but all roughly respond in a few tens of milliseconds, which matches the scale of the membrane time constant of the neurons in the models.

### Temporal Logic

The CRIREL logic gates are not limited to processing the differences between the magnitude of two inputs, but can process the differences in the timing between two inputs [Fig 4a]. Here we define two inputs arriving at the same time as (1, 1). Likewise, two inputs arriving at different times are (1, 0) and (0, 1). As before, the absence of any signal is considered (0, 0). Note that the magnitude of all inputs is the same.

**Figure 4:**
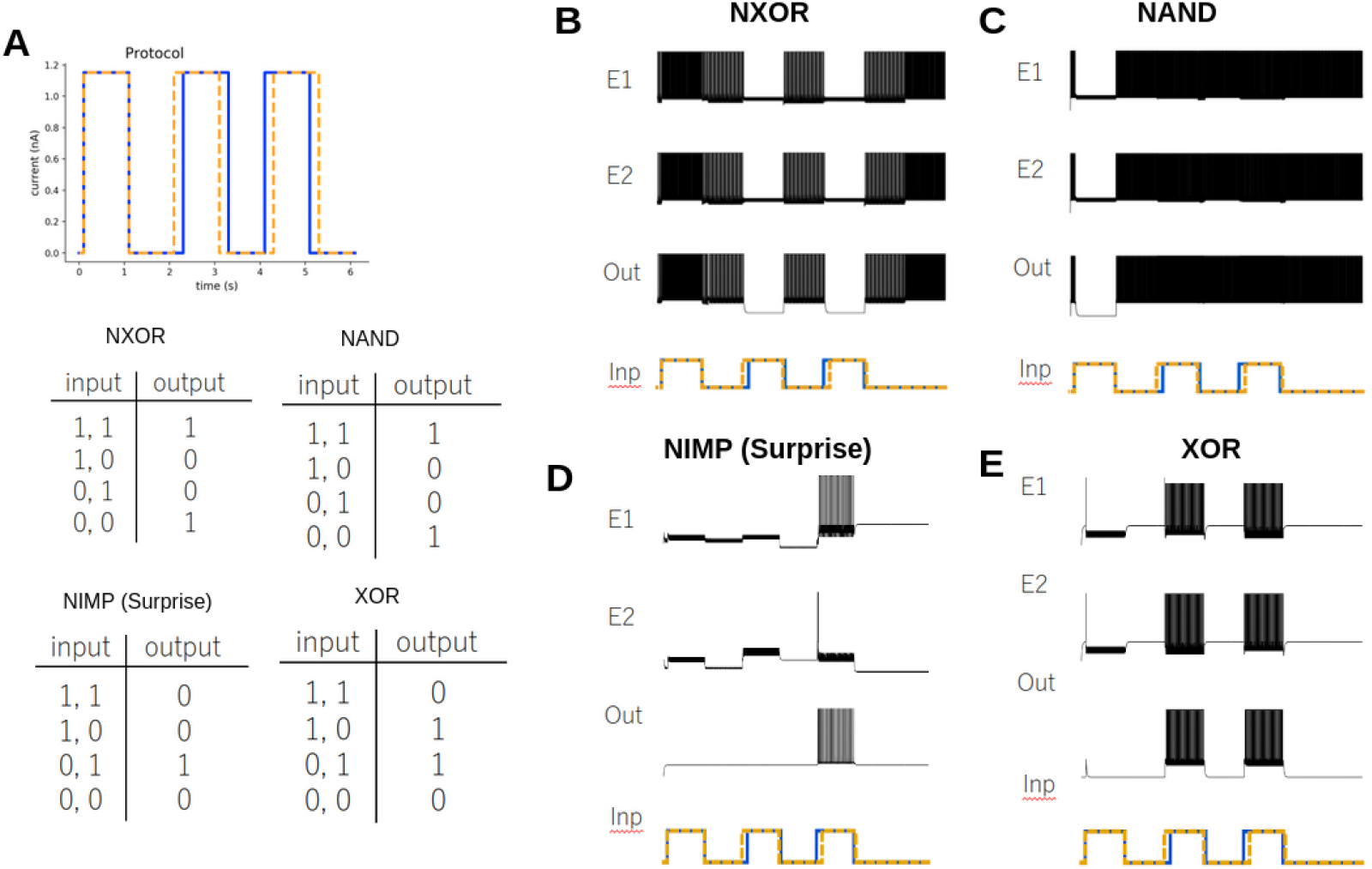
Temporal Logic. A) Protocol for the Temporal Logic Gate. The difference between the timing of the square pulses determines the symbols (1, 1), (1, 0), (0, 1) and (0, 0). The difference in timing is 10 *ms*. The difference between signals is exaggerated for clarity. The Truth tables for NXOR and NAND are shown. B) The LIF version of an NXOR gate. The activity of the excitatory neurons (E1 & E2) and the output neuron (Out) are shown. A cartoon representation of the input protocol is shown. C) The LIF version of a NAND gate. The activity of the excitatory neurons and the output neuron are shown. D) The LIF version of the NIMP gate. This gate is the logical negation of the Imply gate. E) The LIF version of the XOR gate.

To demonstrate, we show an NXOR gate (not exclusive or) [Fig 4b]. Here, the network and thus output neuron fires in the absence of input (0, 0). Whenever the signal (1, 1) is present, the firing rate increases. However, whenever two signals arrive at different times (1, 0) and (0, 1), the circuit turns itself off, and the output neuron is silent. We can easily transform the network into any other of the 8 logic gates. Here, we show the NAND gate [Fig 4c], again the absence of signal (0, 0) is firing, but the network fires when there is a timing difference (1, 0) and (0, 1) and is silent when the timing is the same (1, 1). We also highlight the NIMP gate, which is the logical negation of the Imply gate [Fig 4d]. This gate shows no response if the timing is perfectly synchronized (1, 1) or the blue signal leads (1, 0), however shows a response if the yellow signal leads (0, 1). We refer to this also as a surprise gate, as one can imagine if the blue signal is trying to predict the yellow signal (with some time lag) the network remains silent. However, if the prediction fails, the network turns on to alert to a failure of prediction. Finally, we conclude with an XOR gate example [Fig 4e]. It is worth stressing that these examples coexist with the same synaptic weights, only the bias currents change [Supplementary Table 9].

Again, we show that all 8 unique logic gates exist for at least some set of bias currents, and test this for both LIF neurons [Fig 5a] and for IZH neurons [Fig 5b]. The bias currents used to produce these 8 gates can be found in the supplement[Supplementary Table 9 and Supplementary Table 10]. We provide the parameter sweep for 6 of the logic gates as a 2-dimensional sweep of excitatory subsystem bias, and inhibitory subsystem bias [Fig 5c and 5d]. As before, IMP and NIMP require asymmetric bias currents, they don’t fit neatly into a 2-dimensional sweep. We can test robustness by considering magnitude *A* and timing difference Δ*t* as free parameters (Supplementary Figure 3). It is important to note that the synaptic weights are the same as the synaptic weights in the magnitude example, only the input protocol was changed.

**Figure 5:**
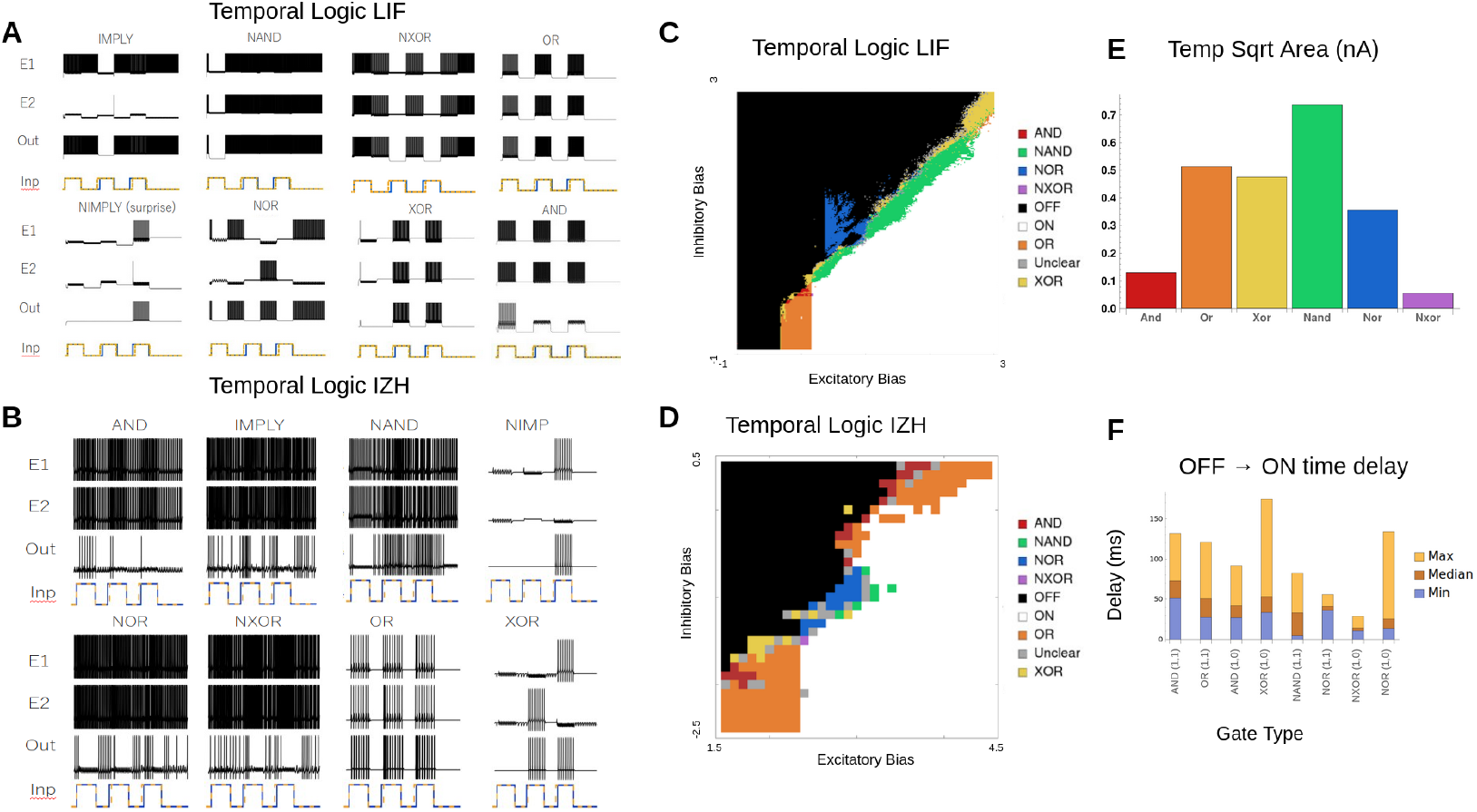
All Temporal gates. A) All 8 Temporal Logic gates for the LIF version of the CRIREL. Both IMP and NIMP have a mirror image that is not shown. B) All 8 Temporal Logic gates for the IZH version of the CRIREL. Both IMP and NIMP have a mirror image that is not shown. C) A sweep through the inhibitory bias and excitatory bias for the LIF model. Each gate is labeled based on its color. D) A sweep through the inhibitory bias and excitatory bias for the IZH model. Each gate is labeled based on its color. E) The robustness of each gate in terms of bias current. It is calculated as the square root of the area for each LIF gate in Panel C. Larger values represent more robust gates. F) The average time from a silent OFF state for the output neuron to the first spike of an ON state for the output neuron. We sampled different biases from the sweep in C.

### Phase Logic

Given the ability to look at the relative timing of two inputs, it is natural to ask if computation can be done on phase differences between two periodically repeating inputs. The input neurons take a pure sinusoidal signal[Fig 6a]. Here it is important to note we do not convert the sine-wave into a spiking signal, rather we feed the raw sine-wave into the CRIREL network directly.

**Figure 6:**
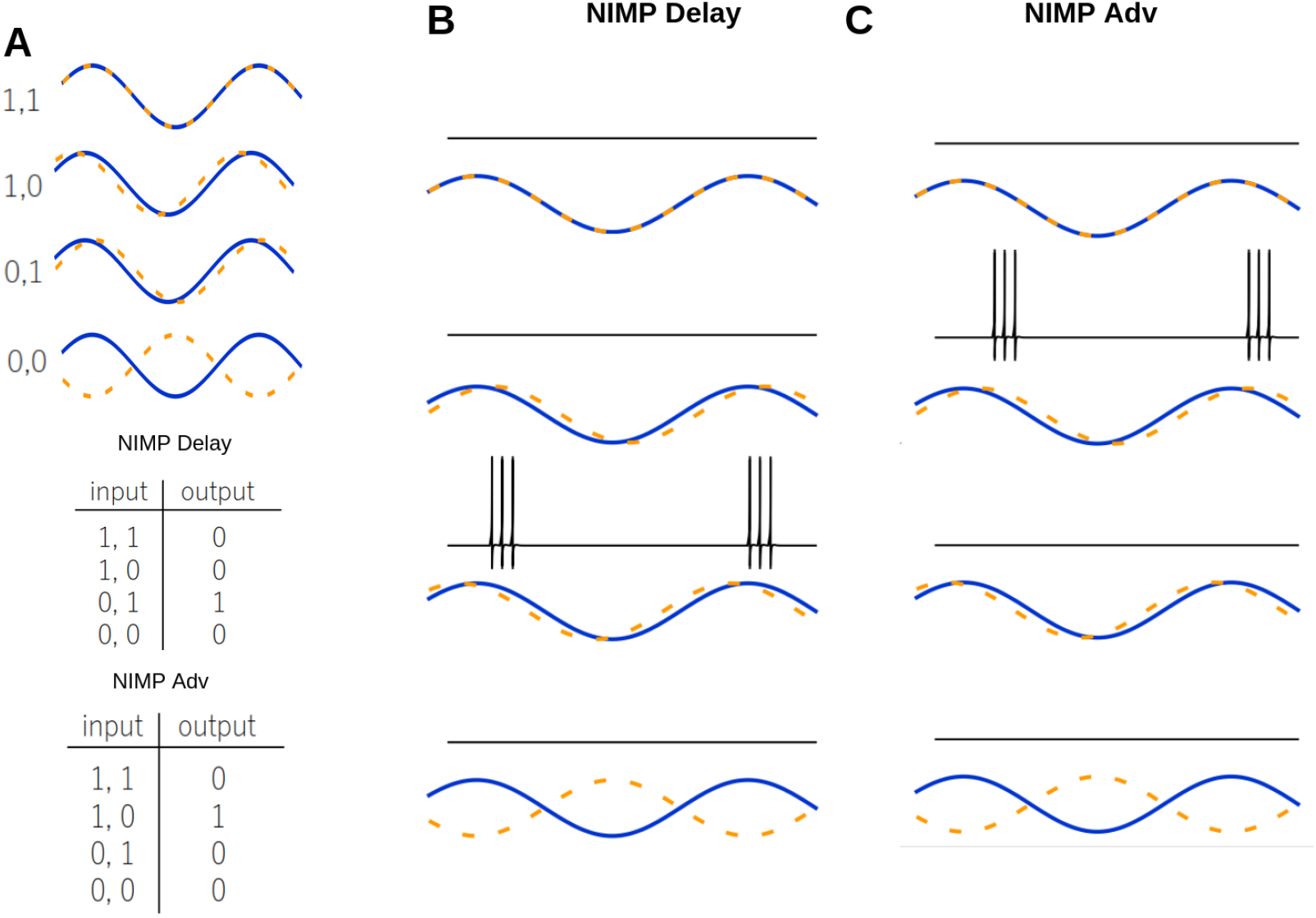
Phase Logic. A) Protocol for the Phase Logic Gate. The difference between the phase of the sine waves determines the input state, ie (1, 1) synchrony, (1, 0) phased advanced, (0, 1) phase delayed and (0, 0) anti-synchronized. The Truth tables for NIMP delayed and NIMP advanced are shown. B) The IZH version of a NIMP delayed gate. The activity of the output neuron is shown. A cartoon representation of the input protocol is shown. C) The IZH version of a NIMP advanced gate. The activity of the excitatory neurons and the output neuron are shown.

As expected, the definition of the inputs and output change. We consider (1, 1) to be when the two input waves are synchronized. Similarly, we consider (0, 1) to be phased delayed and (1, 0) to be phased advanced. Interestingly, we can define (0, 0) as two anti-synchronized inputs. Here, an output neuron is considered ON if, at any time during a full cycle of the sinusoidal input, the output neuron spikes. Often, the output neuron will fire during the maximum of the input wave [Fig 6b and 6c], but for anti-synchronized (0, 0) the output neuron fires during the average value [Fig 7a NOR].

**Figure 7:**
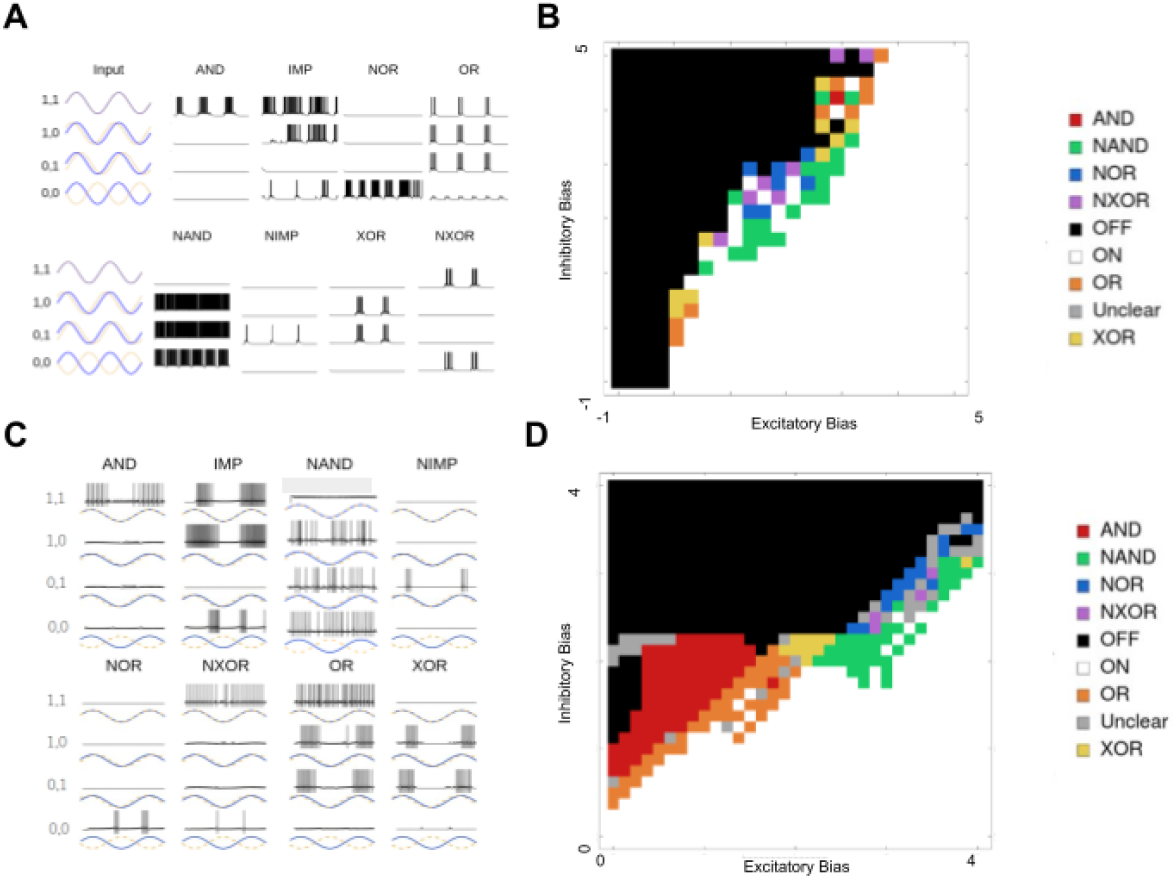
All Phase gates. A) All 8 Phase Logic gates for the LIF version of the CRIREL. Both IMP and NIMP have a mirror image that is not shown. B) A sweep through the inhibitory bias and excitatory bias for the LIF model. Each gate is labeled based on its color. C) All 8 Phase Logic gates for the IZH version of the CRIREL. D) A sweep through the inhibitory bias and excitatory bias for the IZH model. Each gate is labeled based on its color.

As a concrete example, we point out 3 examples: AND [Fig 7a AND], NIMP advanced [Fig 6b] and NIMP delayed [Fig 6c], and NOR[Fig 7a NOR]. For example, AND acts as a coincidence detector and fires only when the two signals are synchronized [Fig 7a AND]. NIMP (or Not Imply) we call the surprise detector[Fig 6b], as if you imagine one wave predicting the other wave, then the output neuron will only fire when the prediction lags behind the measurement, that is the prediction is no longer accurate. More concretely, that means NIMP’s output neuron fires whenever it is in the (1, 0) state, not the (0, 1). Notice that the distinction between (1, 0) and (0, 1) is arbitrary, and as such there are two symmetric versions of NIMP that can be easily switched between by flipping the asymmetry present in the bias currents [Fig 6c]. The NOR gate [Fig 7a] only fires when the two waves are perfectly anti-synchronized. It is silent in the other three cases.

We also show that all 8 unique logic gates exist for phase type computations, and test this for both LIF neurons [Fig. 7a] and IZH neurons [Fig. 7c]. The bias currents used to produce these 8 gates can be found in the supplement (Supplementary Table 11 and Supplementary Table 12). We provide the parameter sweep for 6 of the logic gates as a 2-dimensional sweep of excitatory subsystem bias, and inhibitory subsystem bias for LIF[Fig 7b] and [Fig. 7d]. Because IMP and NIMP require asymmetric bias currents, they don’t fit neatly into a 2-dimensional sweep.

It is also important to discuss how precise the synaptic weights must be. The gates would be useless if they require a specific synaptic weight and cannot tolerate any deviation from this value. We find that most gates are robust up to 10% difference in the synaptic weights [Supplementary Figure 4].

### Application: Adder Circuit

Flexible gate switching is a unique property of CRIREL logic gates. To demonstrate its advantages, we recreated the well-documented 4-bit ripple carry adder circuit, which is a digital logic circuit used for performing multi-bit binary addition. It is composed of four 1-bit full adders [Fig 8a]. A full adder takes in three inputs, the two summands, and the carry from the previous digit [Fig 8b]. This is fed into two CRIRELs, one for computing the sum and the other for computing the carry. We add a buffer neuron to regularize the output firing rate. Note that a traditional transistor-based adder requires 9 NAND logic gates [Fig 8c] [55], while only two neuromorphic logic gates are required.

**Figure 8:**
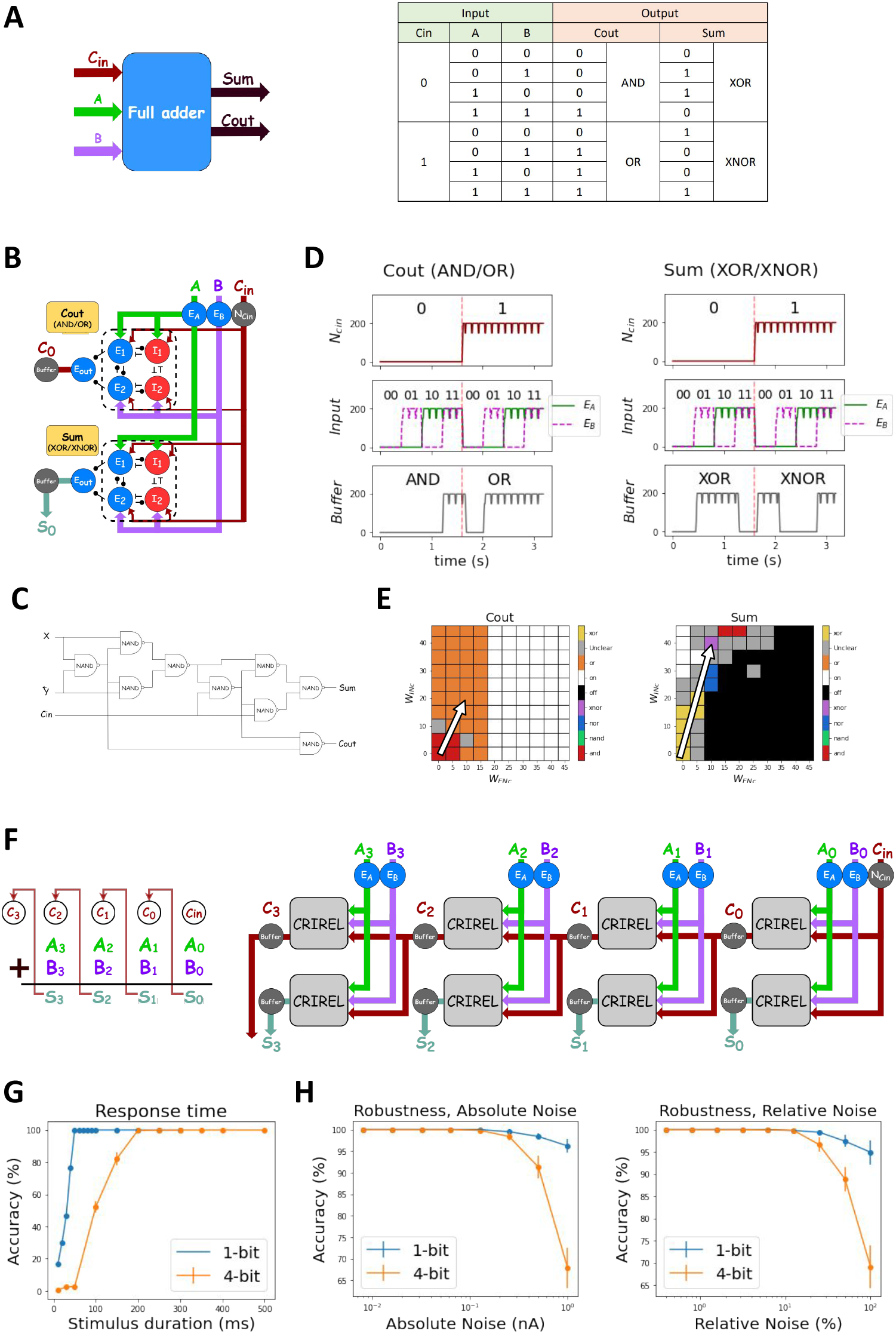
An Adder Application for CRIRELs. (A) The schematic and the truth table of full adder. Carry-out (Cout) toggles between AND and OR gates, while the summation (Sum) toggles between XOR and NXOR gates. (B) The networks of 1-bit full adder. (C) A classical 9-gate 1-bit adder, note that it has 7 more required logic gates than the gate switching adder. (D) The output firing rates of two CRIRELs. (E) Parameter space for the control neurons, when the neuron is off the network switches from the starting gate AND and XOR to NXOR and OR. The synaptic weight of the control neuron is tuned such that the resulting constant current changes the bias current to switch to the correct gate. (F) The schematic and network of 4-bit ripple carry adder (G) The response time of adder. A 1-bit adder requires a response time of 50 *ms* and a 4-bit ripple carry adder needs 200 *ms*. (H) The adder composed of CRIRELs can tolerate absolute noise of 0.1 *nA* and relative noise of 10% in its bias.

Here, it is important to note that the two summands are the traditional magnitude inputs from above. Here, the carry induces a change in the bias current of the CRIRELs to switch the logic gate from AND to OR for the carry CRIREL, and XOR to NXOR for the sum CRIREL [Fig 8d]. The bias currents used to switch can be found in [Supplementary Table 13] and are graphically represented as well [Fig 8e].

It is also worth noting that to ensure output amplitude matches output magnitude, we added a buffer neuron to match the output firing rate (i.e., magnitude) to the input firing rate for AND, OR, XOR, and NXOR logic to work. It works by having a sharp threshold such that it reaches the maximum firing rate with any input from the output neuron. To see the parameter of the buffer neuron see the methods [Supplementary Table 2]. One particularly interesting implication of the full-adder is that CRIREL circuits can be combined together to create cascading logical operations. This is by no means limited to magnitude logic (as with the full adder example), but extends to phase and temporal computation as well [Supplementary Figure 5].

To implement a 4-bit ripple adder, we connect four 1-bit full adders together [Fig 8f]. A 1-bit adder requires at least 50 milliseconds to generate a correct response time [Fig. 8g]. Carry propagation delay necessitates a minimum 200-millisecond delay for a 4-bit ripple carry adder.

We investigated CRIREL’s noise tolerance, which is essential to a network’s usability. We applied Gaussian noise to the bias and measured the accuracy of the logic operation (see Materials and Methods). We quantified both absolute noise (measured in nA) and relative noise (percentage of bias current). Results show CRIREL-based adders have a 0.1nA tolerance for absolute noise and a 10% tolerance for relative noise [Fig. 8h] (NXOR is an exception, ranging between *−*10% to +5%).

### Application: Reconfigurable Motion Detection

Conventional motion detectors such as the Hassenstein-Reichardt model only detect a fixed direction it is set to detect [56]. Our NIMP gates can perform conditional motion detection by ignoring the panning motion and detecting only the moving object in the scene. We demonstrate how NIMP gates can be reconfigured to detect objects moving against optical flow induced by a panning camera. This functionality relies on two NIMP gates that are anti-symmetric counterparts of each other [Fig. 9a]. NIMP gates can be used to detect either rightward or leftward optical flow. For simplicity, we can consider optical flow on a 1 dimensional pixel array[Fig. 9b] Here we have a collection of white pixels moving either rightward or leftward. The circuit compares the pixel value of two adjacent pixels. Depending on the chosen bias currents, the circuit is either rightward-selective or leftward-selective, firing spikes whenever bright objects pass by the two pixels being compared. It achieves selectivity by comparing the phase delay between the left and right pixels [Fig. 9b]. Depending on the bias of the network, the network responds only if the right pixel lags the left pixel, or if the left pixel lags the right pixel. We can now consider a panning camera, again considering a 1-D pixel array. The camera either pans right or left, inducing a flow in the opposite direction. At 1000 milliseconds an object appears moving against the optical flow induced by the panning [Fig. 9c]. The NIMP gate’s directional sensitivity can be tuned to pick up either the object moving against the flow[Fig. 9d left] or the flow itself[Fig. 9d right]. They can be reprogrammed depending on the direction of the panning camera. To detect an object moving with the flow, the directional selectivity is set to be opposite the flow induced by the panning camera. While this is just a simple setup to demonstrate a proof-of-concept, the key observation is that the camera pan can send a signal to the CRIREL circuits and reconfigure the network’s receptive field to be leftward or rightward selective. There is no need to change synaptic weights to change the network’s receptive field.

**Figure 9:**
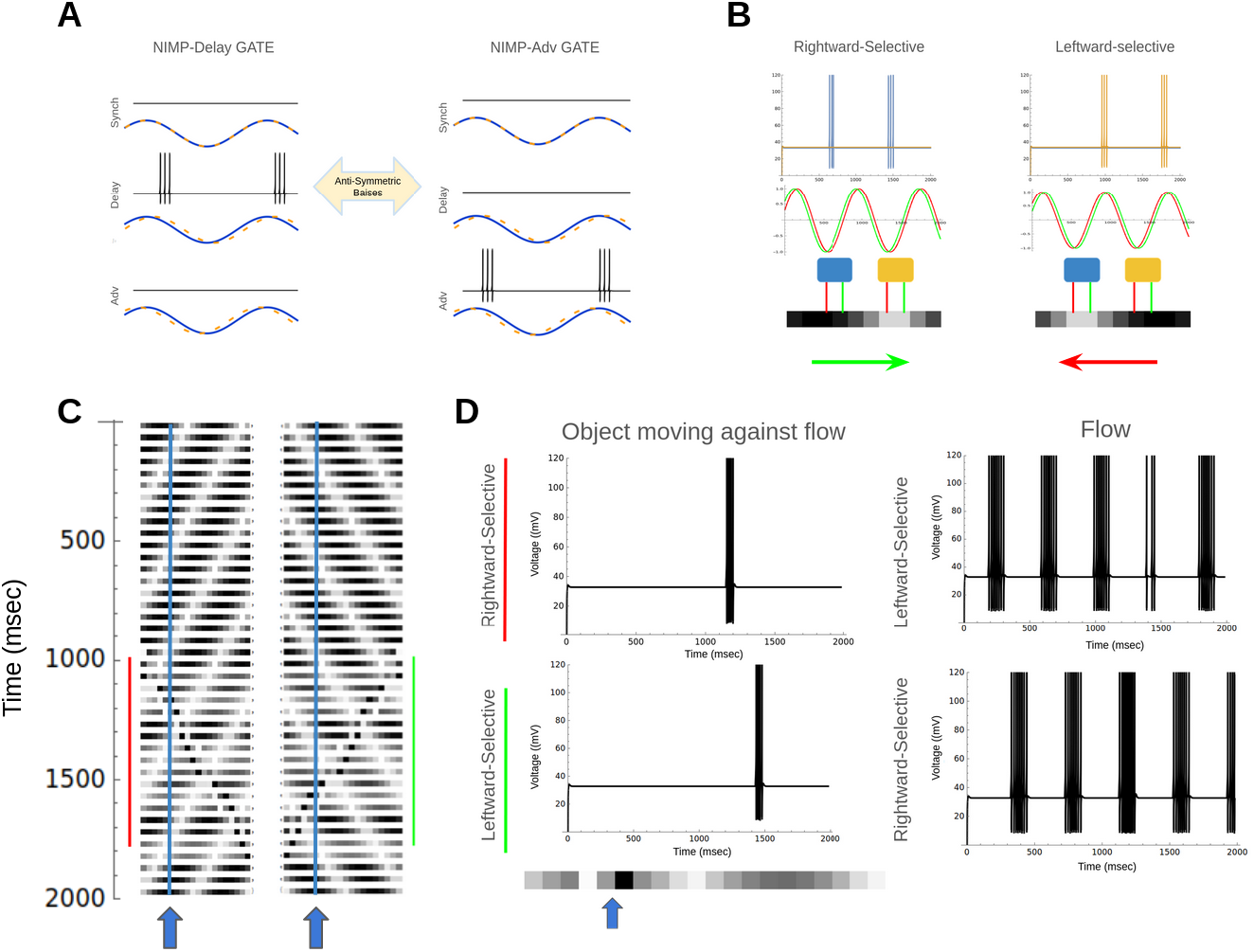
Detecting motion against a moving background A) The Not Imply (NIMP) gate has two antisymmetric versions that can be easily switched between by flipping the bias current asymmetries. B) Two circuits with different antisymmetric bias currents are sensitive to either rightward-moving white pixels or leftward-moving white pixels. C) two examples of a leftward/rightward moving flow with an object moving rightward/leftward at 1000 ms. D)Circuits with NIMP biases comparing two adjacent pixels (indicated by the blue and yellow arrows). The blue circuit detects rightward motion, the yellow circuit detects leftward motion. The network can distinguish between flow and the object moving against the flow.

### Independence of Neuronal Unit

We have stressed that the underlying neuron model that the CRIREL circuit is composed of does not matter. Any neuron with a monotonic saturating IF curve can form a flexible neural network. However, the implementation need not be spiking. Components with saturating non-linearity can also perform flexible logic. Any system of the following form can have flexible and reconfiguable logic.

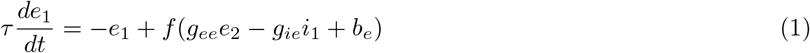

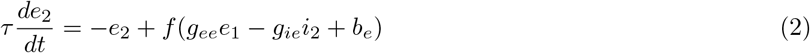

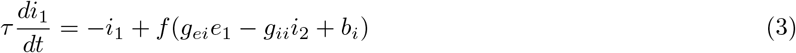

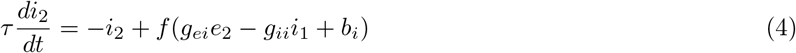

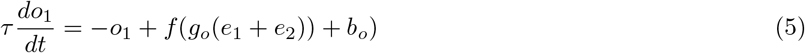

Here we show two implementations of *f* given by a clipped ReLU [Fig. 10a] and a sigmoid function[Fig. 10b].

**Figure 10:**
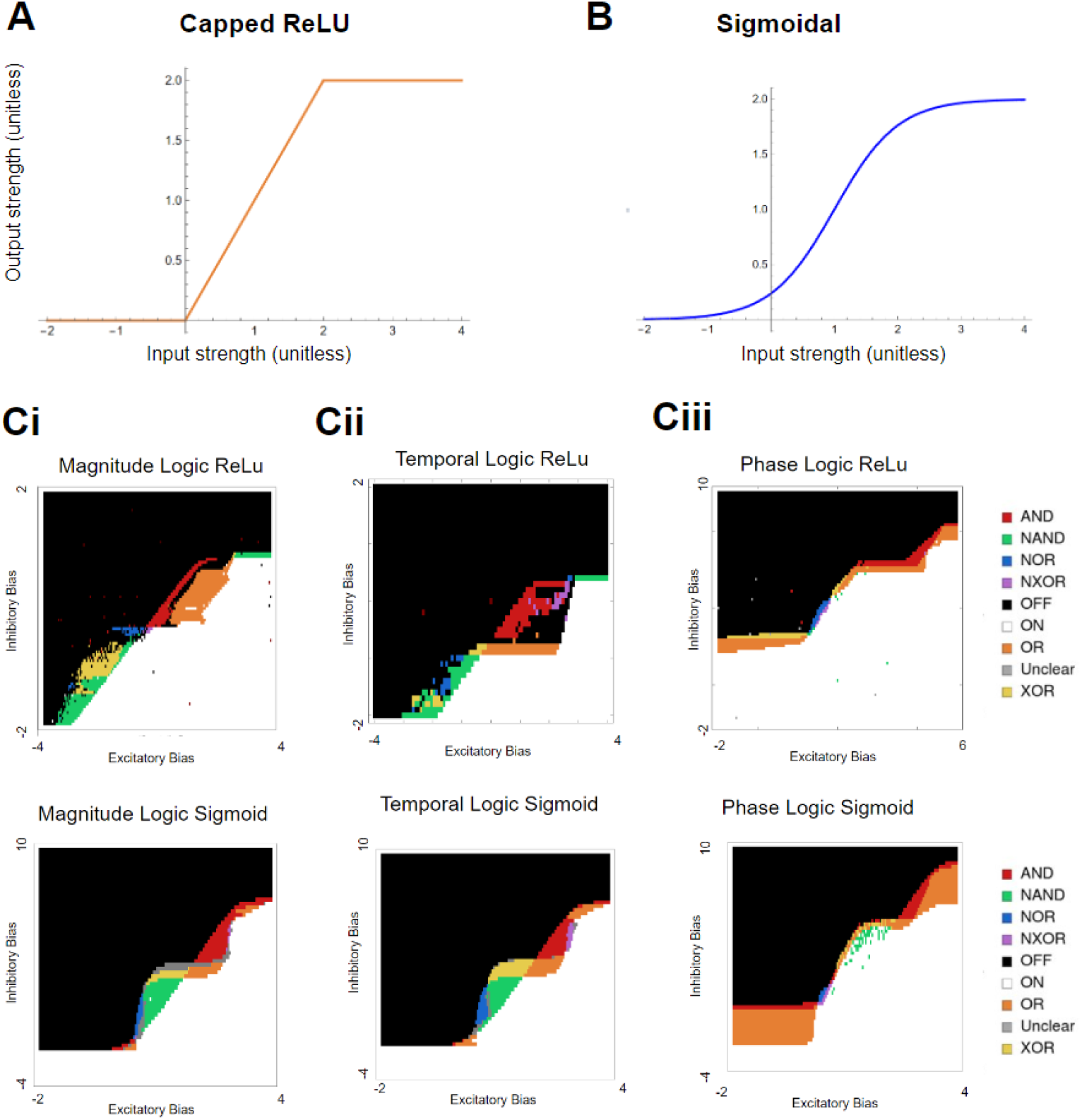
Examples of saturating rate fire neurons in CRIREL circuit. A) Capped ReLU function activation curve. B) Sigmoidal function activation curve. Ci)A sweep through the inhibitory bias and excitatory bias for magnitude computation. Each gate is labeled based on its color. Cii)A sweep through the inhibitory bias and excitatory bias for temporal computation. Ciii) A sweep through the inhibitory bias and excitatory bias for phase computation.

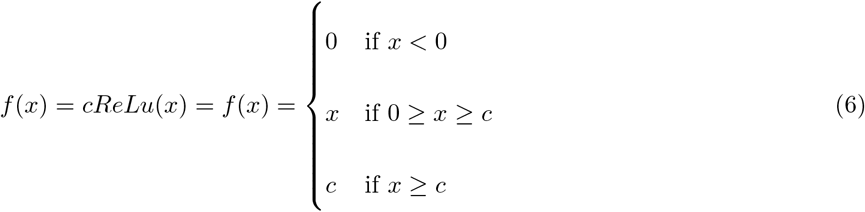

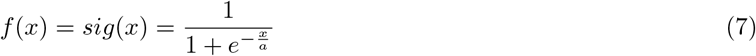

As before, we can test the three different input types: magnitude [Fig 10ci], temporal [Fig 10cii], and phase[Fig 10ciii] for both the sigmoidal function and the clipped ReLU. Again, all 6 symmetric logic gates can be found by sweeping through *b*_*e*_ and *b*_*i*_. It is important to note that there can be oscillatory solutions present. While outside the scope of the paper here, these are useful for pace-making and central pattern generation standpoint (see [9] for a thorough investigation of oscillatory behavior).

## Discussion

In this study, we conclusively show that small spiking recurrent neural microcircuits are incredibly flexible. Here, by picking a set of synaptic weights such that the bias currents can uncontrollably unfold the double-cusp bifurcation, we are able to switch between an incredibly diverse range of different operations. To concretely define unique functions, we used logical truth tables to associate each function with a logic gate. Moreover, we showed that these logic gates can process three different types of inputs, namely, magnitude, temporal, and phase. Furthermore, we found that the gates are robust against membrane noise, and operated over a range of synaptic weights. Strikingly, we showed that these results were independent of the neuron implementation, as we found working versions in LIF, IZH neurons (in the integrator regime), and firing rate models. While we tested the network with all integrator neurons [49], it would indeed be interesting to test the network with the IZH neurons set as resonators [42].

We also investigated two new types of information processing: differences in the arrival of signals, temporal, and differences in the phase of periodic signals. We showed that the flexible gate switching can be used to create a 4-bit ripple carry adder. Additionally, we showed how to define logic with the differences between the signals. Moreover, we showed that this logic can be used to detect events. Specifically, these networks can rapidly and flexibly reconfigure to detect either a rightward traveling white pixels, or a leftward traveling white pixels. Finally, we show that the reconfigurability is a property of the network, not the underlying neuronal model.

This is in line with previous research showing that recurrent neural networks are more flexible than their feedforward counterparts [9]. Specifically, this study takes advantage of a nearby cusp bifurcation [53]. This tracks with the plethora of studies that show networks near bifurcations are inherently more flexible [33, 49, 50, 51, 52, 53]. Studies have been able to embed complex functions into networks that are near multiple bifurcations [33, 38, 48], and other studies have shown, that networks far away from bifurcations means that the network has redundant neurons [53]. Here, we push this idea to its limit by fitting 24 unique functions into a 4-neuron microcircuit. Additionally, we show that one can rapidly switch between these functions without changing the synaptic weights, making use only of the bias currents.

This provides a particularly powerful technique, as often switching functions is seen as a neuromodulatory process [10, 19, 20, 21], or a synaptic weight process [36, 37, 38, 39, 40]. While clearly, these process can and do change a network’s function, they are not the only means. Contextual information and top-down control have been shown to optimize [23, 25, 50, 51, 27, 57, 58], or even switch, the function of decision networks in the prefrontal cortex [13, 59, 60]. Here, we show that bias current (which can be thought of as weak constant input) can rapidly switch the function of a microcircuit, allowing a potential mechanism for integrating contextual information or top-down control. This is a tantalizing direction for future investigation, especially if a learning algorithm can be used to embed multiple tasks into recurrent networks.

Currently, feedforward spiking neural metworks (SNNs) are also capable of computing logic operations [34, 35], with modern implementations operating on single spikes. While these models excel in quick computation due to their single-spike operation, they are more susceptible to noise, as any variation in spike timing can disrupt the logic computation [Supplementary Table 14].

In these models, synaptic weights are precisely tuned for each logic gate, which helps minimize the neuron count and synapse number[Supplementary Table 14] [34]. Despite this, no feedforward network currently can robustly and flexibly adjust to all possible gates by merely changing the bias current. More critically, no feedforward network can compute temporal logic.

This limitation arises because a feedforward network possesses one fixed point and lacks both working memory and temporal dynamics. After the input delay has passed, an input of (1,0) or (0,1) becomes indistinguishable from an input of (1,1). Consequently, distinguishing gates XOR and NXOR from always-off (OFF) and always-on (ON) gates becomes impossible.

While our network here has a particular neuromorphic implementation, we are not limited by these implementations. Our results show that any sigmoidal saturation is capable of generating a flexible and reconfigurable circuit. This leaves the door open to many potential applications. The sigmoidal saturation means reconfigurable electronic circuits could be made with memristors [5, 4, 46] or homojunctions [45]. Moreover, our circuit’s compatibility with neuromorphic architectures could find use in neuromorphic chips [61, 62, 63], especially with liquid state machines [33, 63]. Finally, given that the network was reconstructable with Izhikevich neurons [49] it is conceivable that our neural circuit can be grown in a real brain organoid [7, 8].

Another potential avenue for future work involves the robustness of some of the gates. Gates like AND were the least robust of the operations, being the least tolerable to noise. However, it is known that gap junctions are capable of averaging out the activity levels of two neurons. If a gap junction (which is commonly observed in parvalbumin-containing neurons [64, 65]) is added between the inhibitory neurons, this should expand the basin attraction associated with the (1, 1) state. A larger (1, 1) state should translate into more robustness to noise for the AND state, at the cost of robustness of the XOR state. This remains to be thoroughly tested.

While we tried to be as comprehensive with input processing types, we cannot be exhaustive. Several interesting directions remain for future work. Is there computation based on differences in frequency? Would a network composed of resonators be ideal for detecting differences in frequency? On the other hand, could the order in which symbols are presented (1, 1) then (0, 1) be different than (0, 1) then (1, 1)? Could one leverage this to construct finite-state machines? These are tantalizing open questions.

Finally, while we give a useful account of small microcircuits, the approach needs to be expanded to larger networks. Some studies have successfully embedded arbitrary motor commands into E/I balanced networks near multiple bifurcations [33]. It remains to be seen how many unique functions can be embedded in a larger network, and if this technique of unfolding the bifurcation in predictable ways can be of use. Using our full adder example, we demonstrated how CRIRELs can be interconnected to form cascading logic circuits, where the output of one circuit serves as the input for the next. We showed logic can be cascaded for magnitude (Fig. 8), phase, and temporal computation [Supplementary Figure 5]. However, this is far from a comprehensive list of possible logic circuits. Future work is needed to explore other higher-order circuits composed of flexible CRIRELs. Applications can include motion control [1], motor planning [33], and event-based image processing [66, 67]. Specifically, can learning be used to configure a group of 4-neuron CRIREL circuits, or even a larger, more general network? If so, that could open up new opportunities to minimize neuron count but maximize task number. This could have major consequences for edge-deployed neuromorphic computing.

To conclude, we made use of CRIREL networks near a double cusp bifurcation, and were able to control the unfolding of the bifurcation with the bias currents. We demonstrated that these microcircuits have a rich repertoire of 24 unique functions. Ultimately, this provides a computational foundation for how neural adaptability can occur on timescales much shorter than plasticity. This is a novel way to control a neural circuit, and could lead to new types of computing, especially in the growing field of neuromorphic computing.

## Methods

We make use of two neuron models, the Leaky Integrate-and-Fire model, and the Izhikevich model. This is to show that flexibility is a property of the network, not the underlying neurons. Moreover, it shows the flexibility is robust to model choice.

### Leaky Integrate-and-Fire Model and Synaptic Model

We implement our Leaky Integrate-and-Fire (LIF) model using our in-house simulator Flysim [68]. Recall LIF neuron dynamics are given by the equation

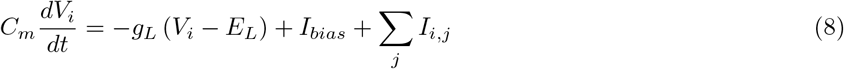

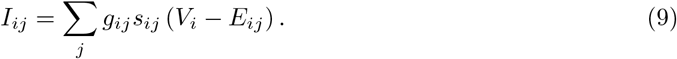

*V*_*i*_ is the membrane potential of the *i*^th^ neuron, *C*_*m*_ is the membrane capacitance, *g*_*L*_ denotes the leak conductance, and *E*_*L*_ represents the leak potential. Here, *g*_*ij*_ corresponds to the synaptic conductance between neurons. We use *s*_*ij*_ to denote the synaptic variable, and *E*_*ij*_ denotes the synaptic reversal potential. Most importantly *I*_*bias*_ is the constant bias current we manipulate to generate the diverse set of logical operations.

We utilize an excitatory synapse (AMPA-like), referred to here as Exc, and an inhibitory synapse (GABA-like), referred to here as Inh, as the neurotransmitters for excitatory and inhibitory neurons. We refer to the two excitatory neurons as E1 and E2, and the two inhibitory neurons as I1 and I2 (see Fig 1A). They are almost always considered symmetric, except when different inputs or bias currents are provided. The equations for the synapses are shown below and parameters for various neurons are given in Supplementary Table 2.

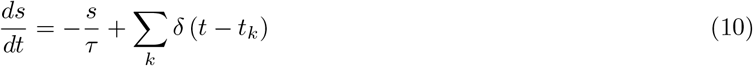

When the membrane potential exceeds the threshold voltage (*V*_*threshold*_), a neuron generates a spike and then is reset to (*V*_*reset*_). There is no change in voltage until after the refractory period lapses.

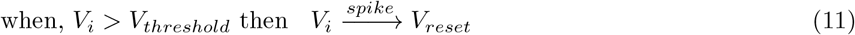

The parameters for the LIF neuron are shown in Supplementary Table 3. We must stress that the synaptic weights for every function are the same. The synaptic weights used for the LIF model are listed in Supplementary Table 4.

### Izhikevich Model

The Izhikevich neuron is a nonlinear generalized quadratic integrate and fire neuron [42, 49]. We implement the Izhikevich neuron (IZH) in Mathematica. We use the following version of the IZH neuron

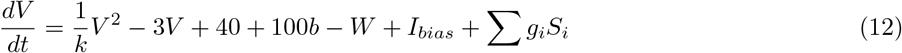

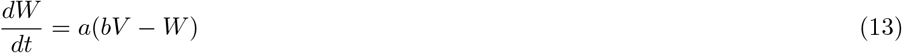

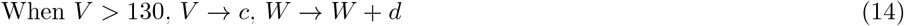

Voltage was set to be akin to millivolts with an always positive voltage, as to aid eventual implementation on a neuromorphic chip. When the voltage hits 130, the system resets to voltage *c* and the gating variable *W* is reset to *W* + *d*. Again, we use *I*_*bias*_ to denote the constant bias current we manipulate to generate the diverse set of logical operations. The synapse gating variables *S* were governed by the following equation

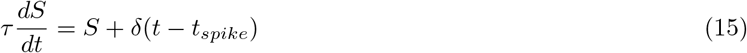

where *τ* is a time constant, and *δ* is the Dirac delta function, raising the value of the synaptic gating variable *S* by one unit every spike. The parameters (which should be considered unitless) are given in Supplementary Table 5. We choose the parameters such that the neuron is an integrator.

Synaptic Weights between each neuron are given in Supplementary Table 6.

### Protocol for Binary Inputs

To properly define a unique function, we use logic gates as a useful classification of different functions. Classically, the input of a logic gate operates on a binary input of two digits, either 0 and 1. There are 4 possible combinations of input (1, 1), (0, 1), (1, 0), and (0, 0) and two possible outputs, 1 or 0. There are 16 possible combinations of inputs and outputs [Supplementary Table 1].

However, a neural circuit is not constrained to be binary. Neurons can respond to many different properties of an input. Here, we define an (1, 1) input as two inputs that are approximately the same (see supplemental material for the precise range of sameness). If one input is larger in magnitude, it is (1, 0) or (0, 1). If one input arrives before the other (so-called temporal logic), we can also define it as (1, 0) or (0, 1). Finally, if one input is phase delayed to the other input, we can classify it as (1, 0) or (0, 1). For both magnitude and temporal logic, (0, 0) is defined as an absence of input. However, for phase logic, (0, 0) is defined as perfect asynchrony. The truth tables can be found in Supplementary Table 1.

In the LIF implementation, for both magnitude logic and temporal logic, we present the input neurons of the network with two square inputs. For the 1 symbol, the square wave has an amplitude of 5 nA. The 0 symbol has an amplitude of 3.5 nA (except during robustness testing).

In the IZH implementation, for both magnitude logic and temporal logic, we directly present the CRIREL circuit with the square pulse. For the 1 symbol, the square wave has an amplitude of 2.5. The 0 symbol has an amplitude of 2.5 (except during robustness testing). Recall that the Izhikevich neuron is unitless.

For phase logic, in both versions, we present a sinusoidal input directly to the CRIREL circuit. In the LIF model, the wave has a period of 40 ms, and an amplitude of 2 nA. Phase delay was defined as 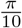 radians. For the IZH model, the wave has a period of 400 ms, and an amplitude of 1. Phase delay was defined as 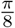 radians.

### Categorizing Logic Gates

To categorize the logic gates for the two neuromorphic implementations, we used a protocol in which the input given to the circuit follows the following order: (0, 1), (1, 1), (1, 0), (0, 1), with a duration of 1 time units for each input, and a window of 0.5 time units between each input. We then check whether the output neuron has any spikes. If so, we label that output as 1. If not, it is 0. We then construct a logic gate truth table and assign a logic gate accordingly

To categorize the non-neuromorphic implementations, each pulse was 5 time units long, with a 5 time unit pause between the inputs. We took the value of the output neuron in the middle of the pulse (2.5 time units after input onset). If it was above the threshold of 1.5 units, we considered it 1 else 0. For phase, we considered the circuit on if the maximum value ever exceeded a threshold of 1.5 units for the 40 time unit simulation.

### Accuracy Testing for Ripple Carry Adder

To test the accuracy and robustness against noise, we inject all neurons in the 4-neuron circuit (E1, E2, I1, I2) with a Gaussian white noise. The mean of the white noise is always zero. We sweep exponentially through increasing standard deviation strength. For the absolute noise, we sweep from a standard deviation of .01 *nA* to a standard deviation of 1.0 *nA*. For the relative noise condition, we sweep from a standard deviation of 0.1% of the maximal bias of each neuron to 100% of the maximal bias of each neuron. We present 1000 randomly chosen pairs of 4-bit numbers, and run 10 trials (each a different Gaussian noise). If the adder returns the correct result for all pairs of numbers and trials -that is 100 % accuracy, we consider it robust.

### Directional Selectivity

To test the phase NIMP logic gate’s ability to flexibly respond to directional stimuli, we build a 1-D array 20 pixels long. We generate a sinusoidal traveling wave within this 1-D pixel array. The luminosity *p* of each pixel is a sinusoidal function of pixel location *x* and *t* in milliseconds. The equation is given by

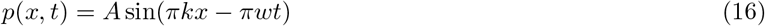

with 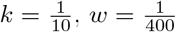, and *A* = 1.0. Each CRIREL circuit is connected to adjacent pixels *x* and *x* + 1, such that input currents *in*_1_(*t*) = *p*(*x, t*) (red) and *in*_2_(*t*) = *p*(*x* + 1, *t*) (green). As this is just proof-of-concept, we only plot the CRIREL circuit at pixel *x* = 10 and 11 (blue) as well as *x* = 15 and *x* = 16 (yellow).

In order to generate NIMP Logic in the phase mode, we used the Izhikevich version of the phase logic. We used the bias currents in the Supplementary Table 12 for both leftward and rightward detection. Note that the different direction selectivities are anti-symmetric. This allows a CRIREL sensitive to leftward motion to be reconfigured to rightward detection simply by changing the bias currents.

To challenge the network more, we generate an “object” moving against the sinusoidal traveling wave. To implement this, we use the following equation for pixel luminosity

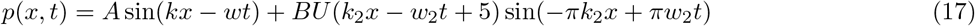

Here, *U* is the Unit Box function with *U* (*y*) = 1 if 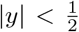 else 0. We left *A, k*, and *w* the same as above. We set *B* = *−*4, 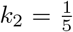 and 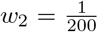. This generates a 5-pixel wide object moving at the opposite speed to the traveling wave.

In order to detect an object moving against the flow, we use a single CRIREL circuit in our array. For demonstration purposes, we put the CRIREL circuit at pixels 5 and 6. Depending on the bias current chosen, the CRIREL circuit is sensitive to leftward or rightward motion. If the panning camera is moving leftward (inducing a rightward flow), the CRIREL circuit can be set leftward to detect an object moving against the flow, or it can be set to rightward detection to detect the flow. The CRIREL can be reconfigured to do whichever computation the user desires.

### Firing rate models

The network structure is given as

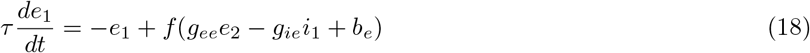

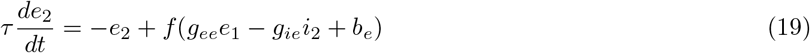

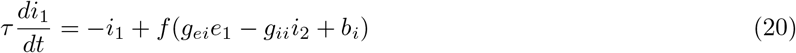

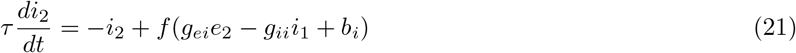

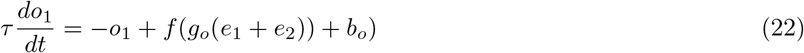

Where *e*_1_ and *e*_2_ are state variables akin to the excitatory neurons. Likewise, *i*_1_ and *i*_2_ are state variables akin to the inhibitory neurons Here we set *b*_*o*_ = 0. Here *b*_*e*_ and *b*_*i*_ are the bias currents we manipulate to change gates. We set *τ* = .25 We offset the initial conditions from zero. We set *e*_1_(0) = .1,*e*_2_(0) = .1, *i*_1_(0) = .02,*i*_2_(0) = .02

The activation function for the sigmoidal version was

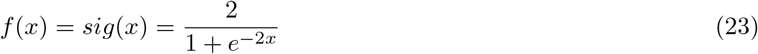

We set the “synaptic” weights as *g*_*ee*_ = 1.5, *g*_*ii*_ = *g*_*ei*_ = *g*_*ie*_ = 2 and *g*_*o*_ = 1.

The activation for the capped ReLU function was

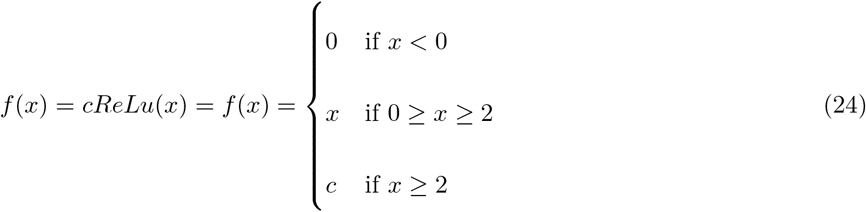

*c* if *x ≥* 2 We set the “synaptic” weights as *g*_*ee*_ = *g*_*ii*_ = 2, *g*_*ei*_ = *g*_*ie*_ = 1.75, and *g*_*o*_ = 1.

## 0.1 Code accessibility

The full source code and parameter files are available at https://github.com/ajw131/CRIREL_Logic_gates [69].

## Declaration of Conflicts of Interest

AJW, BL and CCL are inventors in Taiwan Patent No. #I725914 and United States Patent No. #US12079709 B2 related to the research presented in this paper. Additionally, AJW, BL, and CCL have received royalty payments from Egis Technology Inc. for granting an exclusive license of these patents. The licensing arrangement has been reviewed and approved by The National Tsing Hua University in accordance with its policy on the management and commercialization of intellectual properties. All other authors declare no competing interest.

## Acknowledgements

We would like to thank Dr. Chou P. Hung, and Dr. Andre V. Harrison for feedback on the manuscript. This work is partially supported by the Featured Areas Research Center Program within the framework of the Higher Education Sprout Project, a joint fund from the Ministry of Education (MOE) and the Ministry of Science and Technology (MOST) in Taiwan, 112-2321-B-002-025. This work was also partially supported by National Science and Technology Council (Taiwan), 111-2311-B-007-011-MY3.

## Author Contributions

CCL/KAW provided supervision for the entire project. BL/AJW discovered the logic gates and developed the theory. BL/AJW/CCL developed simulations and robustness tests. BL/MJH/AJW ran simulations. MJH developed the ripple adder. MFC provided critical advice that guided the development of the motion detector and generalized firing rate simulations. AJW/CCL wrote the manuscript. All Authors have approved of the manuscript.

## Notes

### Summary of Updates

A new application was added that shows motion detection. We also showed that the method works in non spiking recurrent networks.

