## Supplemental figures and tables for "Hyper Flexible Neural Networks Rapidly Switch between Logic Operations in a Compact Four Neuron Circuit"

### 1 Supplementary Tables

#### Logic Table

| **AND** | | | **OR** | | | **NAND** | | | **XOR** | | |
|---|---|---|---|---|---|---|---|---|---|---|---|
| **A** | **B** | **Out** | **A** | **B** | **Out** | **A** | **B** | **Out** | **A** | **B** | **Out** |
| 0 | 0 | 0 | 0 | 0 | 0 | 0 | 0 | 1 | 0 | 0 | 0 |
| 0 | 1 | 0 | 0 | 1 | 1 | 0 | 1 | 1 | 0 | 1 | 1 |
| 1 | 0 | 0 | 1 | 0 | 1 | 1 | 0 | 1 | 1 | 0 | 1 |
| 1 | 1 | 1 | 1 | 1 | 1 | 1 | 1 | 0 | 1 | 1 | 0 |
| **NOR** | | | **XNOR** | | | **1 IMP 2** | | | **2 IMP 1** | | |
| **A** | **B** | **Out** | **A** | **B** | **Out** | **A** | **B** | **Out** | **A** | **B** | **Out** |
| 0 | 0 | 1 | 0 | 0 | 1 | 0 | 0 | 1 | 0 | 0 | 1 |
| 0 | 1 | 0 | 0 | 1 | 0 | 0 | 1 | 1 | 0 | 1 | 0 |
| 1 | 0 | 0 | 1 | 0 | 0 | 1 | 0 | 0 | 1 | 0 | 1 |
| 1 | 1 | 0 | 1 | 1 | 1 | 1 | 1 | 1 | 1 | 1 | 1 |
| **ON** | | | **OFF** | | | **1 NIMP 2** | | | **2 NIMP 1** | | |
| **A** | **B** | **Out** | **A** | **B** | **Out** | **A** | **B** | **Out** | **A** | **B** | **Out** |
| 0 | 0 | 1 | 0 | 0 | 0 | 0 | 0 | 0 | 0 | 0 | 0 |
| 0 | 1 | 1 | 0 | 1 | 0 | 0 | 1 | 0 | 0 | 1 | 1 |
| 1 | 0 | 1 | 1 | 0 | 0 | 1 | 0 | 1 | 1 | 0 | 0 |
| 1 | 1 | 1 | 1 | 1 | 0 | 1 | 1 | 0 | 1 | 1 | 0 |
| **A** | | | **B** | | | **NOT A** | | | **NOT B** | | |
| **A** | **B** | **Out** | **A** | **B** | **Out** | **A** | **B** | **Out** | **A** | **B** | **Out** |
| 0 | 0 | 0 | 0 | 0 | 0 | 0 | 0 | 1 | 0 | 0 | 1 |
| 0 | 1 | 0 | 0 | 1 | 1 | 0 | 1 | 1 | 0 | 1 | 0 |
| 1 | 0 | 1 | 1 | 0 | 0 | 1 | 0 | 0 | 1 | 0 | 1 |
| 1 | 1 | 1 | 1 | 1 | 1 | 1 | 1 | 0 | 1 | 1 | 0 |

Table 1: The truth table for all 16 classical logic gates. Six are trivial: always off (false), always on (true), A, B, Not A and Not B gates. For the remaining ten gates, six are symmetric and four are asymmetric. Here we consider AND, OR, XOR, XNOR, NOR, and NAND as the six unique non-trivial symmetric gates. NIMP and IMP constitute the four unique non-trivial asymmetric gates.

#### 1.1 LIF neuron model parameters

| Neuron | Synapse Type | $E_{ij}$ (mV) | $\tau$ (ms) |
|---|---|---|---|
| $E_A$, $E_B$, E1, E2, $E_{Out}$ | Exc | 0 | 2 |
| Buffer | Exc | 0 | 20 |
| I1, I2 | Inh | -90 | 5 |

Table 2: Reversal potentials and time constants for different neurotransmitters in the LIF model.

| Parameter | E1, E2, I1, I2, | $E_{Out}$, $E_A$, $E_B$ | Buffer |
|---|---|---|---|
| $C_m$ $(nF)$ | 0.5 | 0.5 | 0.5 |
| $g_L$ $(\mu S)$ | 25 | 25 | 25 |
| $E_L$ (mV) | -70 | -70 | -70 |
| $V_{\text{threshold}}$ (mV) | -50 | -50 | -50 |
| $V_{\text{reset}}$ (mV) | -55 | -55 | -50.1 |
| Refractory period (ms) | 2 | 5 | 5 |

Table 3: Neurons parameters for the LIF model. Note that only the adder uses the buffer neuron.

| | |
|---|---|
| $g_{ee}$ | 30 |
| $g_{ei}$ | 30 |
| $g_{ie}$ | -60 |
| $g_{ii}$ | -30 |
| $g_o$ | 50 |

Table 4: Synaptic values used throughout the paper in the LIF model, except where explicitly noted.

#### 1.2 IZH neuron model parameters

| Parameter | E1, E2, I1, I2, $E_{Out}$ |
|---|---|
| $k$ | 25 |
| $a$ | $\frac{1}{50}$ |
| $b$ | .2 |
| $c$ | 10 |
| $d$ | 0.1 |
| $\tau$ | 10 |

Table 5: Neurons parameters for the IZH model used in the study.

| | |
|---|---|
| $g_{ee}$ | 3 |
| $g_{ei}$ | 5 |
| $g_{ie}$ | -10 |
| $g_{ii}$ | -10 |
| $g_o$ | 3 |

Table 6: Synaptic values used throughout the paper in the IZH model, except where explicitly noted

#### Supplementary Table Magnitude

| Parameter for Magnitude Logic Gates (LIF) | | | | | |
|---|---|---|---|---|---|
| Logic Gate | $b_{E_1}$ | $b_{E_2}$ | $b_{I_1}$ | $b_{I_2}$ | $b_{out}$ |
| AND | 0 | 0 | -0.2 | -0.2 | 0 |
| NAND | 0.75 | 0.75 | 1.7 | 1.7 | 0.5 |
| OR | 0.2 | 0.2 | -0.5 | -0.5 | 0.5 |
| NOR | 0.65 | 0.65 | 2.1 | 2.1 | 0.5 |
| XOR | 0.3 | 0.3 | 1.4 | 1.4 | 0.5 |
| XNOR | 0.9 | 0.9 | 0.5 | 0.5 | -0.7 |
| IMP | 1 | 0.7 | 0.8 | 1 | 0 |
| NIMP | 0.2 | 0 | 0.85 | 0.85 | 0 |

Table 7

| Parameter for Magnitude Logic Gates (IZH) | | | | | |
|---|---|---|---|---|---|
| Logic Gate | $b_{E_1}$ | $b_{E_2}$ | $b_{I_1}$ | $b_{I_2}$ | $b_{out}$ |
| AND | 1.25 | 1.25 | -1 | -1 | -3.5 |
| NAND | 3.5 | 3.5 | 1.0 | 1.0 | -3.0 |
| OR | 1.0 | 1.0 | -1.0 | -1.0 | 3.0 |
| NOR | 5.05 | 5.05 | 2.6 | 2.6 | -5.0 |
| XOR | 1.55 | 1.55 | 0 | 0 | -2.1 |
| XNOR | 3.0 | 3.0 | -1.0 | -1.0 | -7.25 |
| IMP | 3.6 | 3.7 | -.75 | -.7 | -7.22 |
| NIMP | 1.5 | 0.5 | 1 | 0.5 | -1.5 |

Table 8

#### Supplementary Table Temporal

| Parameter for Temporal Logic Gates (LIF) | | | | | |
|---|---|---|---|---|---|
| Logic Gate | $b_{E_1}$ | $b_{E_2}$ | $b_{I_1}$ | $b_{I_2}$ | $b_{out}$ |
| AND | 0 | 0 | 0 | 0 | 0 |
| NAND | 1.5 | 1.5 | 1.0 | 1.0 | 0 |
| OR | 0 | 0 | -0.2 | -0.2 | 0 |
| NOR | 0.9 | 0.9 | 1.25 | 1.25 | 0 |
| XOR | 0 | 0 | 0.85 | 0.85 | 0 |
| XNOR | 1.25 | 1.25 | 1.15 | 1.15 | 0 |
| IMP | 1.2 | 0.7 | 0.9 | 1 | 0 |
| NIMP | 0.5 | 0 | 0.85 | 0.85 | 0 |

Table 9

| Parameter for Temporal Logic Gates (IZH) | | | | | |
|---|---|---|---|---|---|
| Logic Gate | $b_{E_1}$ | $b_{E_2}$ | $b_{I_1}$ | $b_{I_2}$ | $b_{out}$ |
| AND | 3.35 | 3.35 | 0.35 | 0.35 | -5.45 |
| NAND | 3.5 | 3.5 | 1.0 | 1.0 | -3.6 |
| OR | 1.0 | 1.0 | -1.0 | -1.0 | 3.0 |
| NOR | 4.0 | 4.0 | -0.35 | 0.35 | -7.65 |
| XOR | 0.5 | 0.5 | 2.0 | 2.0 | 3.0 |
| XNOR | 3.75 | 3.75 | -1.0 | -1.0 | -8.0 |
| IMP | 3.6 | 3.7 | -.75 | -.7 | -7.22 |
| NIMP | 0.5 | -0.5 | 1.0 | 1.1 | 2 |

Table 10

#### 1.3 Supplementary Table Phase

| Parameter for Phase Logic Gates (LIF) | | | | | |
|---|---|---|---|---|---|
| Logic Gate | $b_{E_1}$ | $b_{E_2}$ | $b_{I_1}$ | $b_{I_2}$ | $b_{out}$ |
| AND | 4.25 | 4.25 | 3.0 | 3.0 | 0 |
| NAND | 2.25 | 2.25 | 1.5 | 1.5 | 0 |
| OR | 1.0 | 1.0 | 0.0 | 0.0 | 0 |
| NOR | 2.75 | 2.75 | 1.5 | 1.5 | 0 |
| XOR | 1.5 | 1.5 | 0.0 | 0.0 | 0 |
| XNOR | 3.25 | 3.25 | 1.75 | 1.75 | 0 |
| IMP | 2.75 | 3.3 | 1.75 | 1.75 | 0 |
| NIMP | 1.5 | 2.1 | -0.25 | -0.25 | 0 |

Table 11

#### Supplementary Table Phase (IZH)

| Parameter for Phase Logic Gates (IZH) | | | | | |
|---|---|---|---|---|---|
| Logic Gate | $b_{E_1}$ | $b_{E_2}$ | $b_{I_1}$ | $b_{I_2}$ | $b_{out}$ |
| AND | 3.5 | 3.5 | 3.1 | 3.1 | -2.1 |
| NAND | 5.05 | 5.05 | 2.6 | 2.6 | -5 |
| OR | 4 | 4 | 3.5 | 3.5 | -2.1 |
| NOR | 4 | 4 | 4.5 | 4.5 | -2 |
| XOR | 2 | 2 | 2.5 | 2.5 | 1.1 |
| XNOR | 3.6 | 3.6 | 3.3 | 3.3 | -2.05 |
| IMP | 2.99 | 3.13 | 3.4 | 3.5 | 0 |
| NIMP | 0 | 1.5 | 2.5 | 2.5 | 2 |

Table 12

#### Parameters for Full Adder

| Gate | $W_{enc}$ | $W_{inc}$ |
|---|---|---|
| AND | 0 | 0 |
| OR | 10 | 20 |
| XOR | 0 | 0 |
| NXOR | 10 | 40 |

Table 13: Weights from the carry neuron that change the bias of the logic gates to perform addition in the full adder.

#### Comparison to past work

| | Gate | CRIREL | A.A.M. *et al.*[?] | L.M. *et al.*[?] |
|---|---|---|---|---|
| Neuron Number | AND | 5 | 2 | 10 |
| | OR | 5 | 1 | 10 |
| | XOR | 5 | 4 | 10 |
| | NAND | 5 | 5* | 10 |
| | NOR | 5 | 4* | 10 |
| | NXOR | 5 | 7* | 10 |
| | Full Adder | 12 | 13* | 36 |
| Synapse Number | AND | 10 | 5 | 16 |
| | OR | 10 | 2 | 16 |
| | XOR | 10 | 6 | 16 |
| | NAND | 10 | 9* | 16 |
| | NOR | 10 | 6* | 16 |
| | NXOR | 10 | 10* | 16 |
| | Full Adder | 22 | 28 * | 80 |
| Latency (Lower better) | AND | Mag ($\approx$ 25 ms) | 2 ms | $>$ 1 ms |
| | OR | Mag ($\approx$ 25 ms) | 1 ms | $>$ 1 ms |
| | XOR | Mag ($\approx$ 25 ms) | 2 ms | $>$ 1 ms |
| | NAND | Mag ($\approx$ 5 ms) | 3* ms | $>$ 1 ms |
| | NOR | Mag ($\approx$ 10 ms) | 2* ms | $>$ 1 ms |
| | NXOR | Mag ($\approx$ 10 ms) | 3* ms | $>$ 1 ms |
| | Full Adder | Mag ($\approx$ 50 ms) | Not Tested | $\approx$ 1 ms |
| Noise Tolerance (timing) (Higher better) | AND | $\approx$ 80 ms | $\approx$ 2 ms | Not Tested |
| | OR | $\approx$ 80 ms | $\approx$ 2 ms | Not Tested |
| | XOR | $\approx$ 80 ms | $\approx$ 2 ms | Not Tested |
| | NAND | $\approx$ 80 ms | Not Tested | Not Tested |
| | NOR | $\approx$ 80 ms | Not Tested | Not Tested |
| | NXOR | $\approx$ 80 ms | Not Tested | Not Tested |
| Flexible | All Gates | Full | NO | NO |
| Temporal | All Gates | Full | NO | NO |

Table 14: Comparison of the 6 symmetric logic gates between us and two feed-forward (no recurrence) SSNs by Alvaro Ayuso-Martinez *et al.* and Lingfei Mo *et al.* It is important to note that their SSN models are spike based logic operating on a single spike, while CRIREL integrates over multiple spikes. The feedforward models have lower latency, and fewer neurons. Here, $*$ represents unreported gates that can be constructed making use of gates presented in Alvaro Ayuso-Martinex *et al.*[?] but were not implemented. We should note while single spike computations are faster, CRIREL, has better noise tolerance, solely because it requires longer inputs to come to a "decision". Most importantly, feedforward networks report very little flexibility. Moreover, Feedforward networks cannot process temporal inputs as they lack basins of attraction and hysteresis.

#### 2 supplemental fig

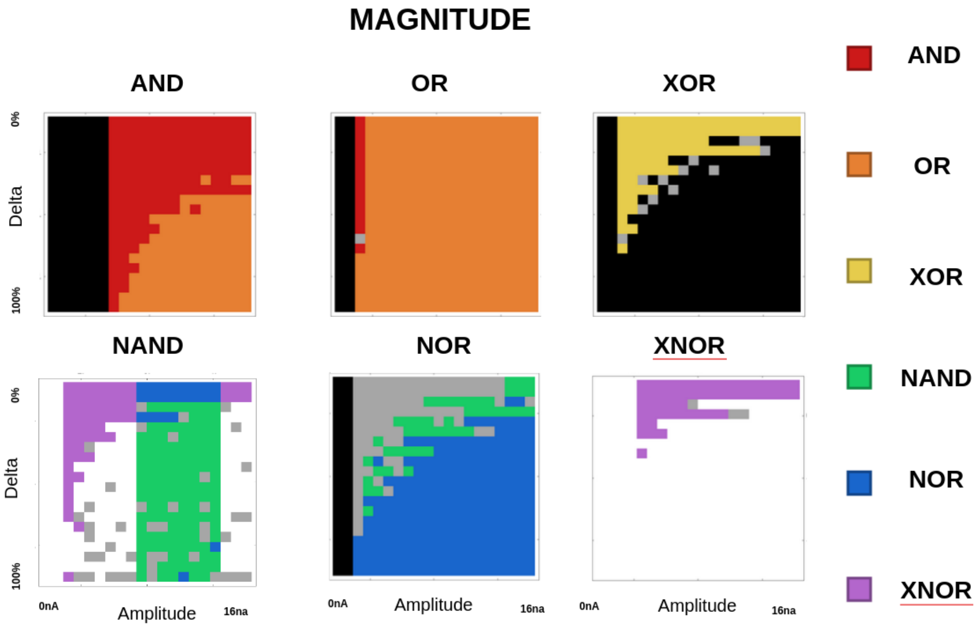


Supplementary Figure 1. A graph of robustness for the 6 symmetric magnitude logic gates as a function of amplitude and the difference in amplitude. Most gates are very robust, except for XNOR.

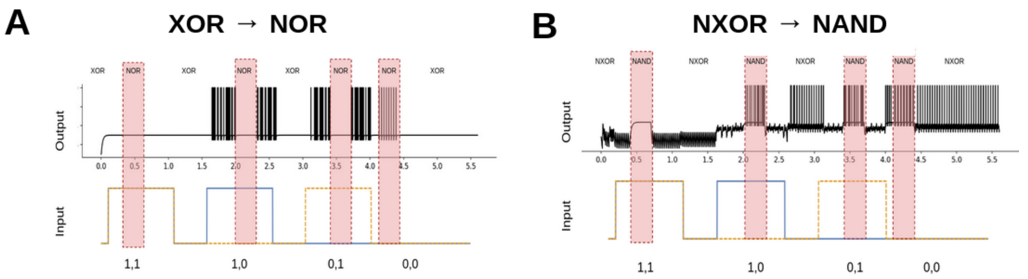


**C** **Robust Gate Switching Possible**

| | AND | IMP | NAND | NIMP | NOR | NXOR | OR | XOR |
|---|---|---|---|---|---|---|---|---|
| AND | | O | O | O | X | X | O | O |
| IMP | O | | O | O | O | X | O | X |
| NAND | O | O | | O | O | O | O | O |
| NIMP | O | O | O | | O | O | O | O |
| NOR | X | O | O | O | | O | X | O |
| NXOR | O | O | O | O | O | | X | O |
| OR | O | O | O | O | X | X | | O |
| XOR | O | O | X | O | O | O | O | |

Supplementary Figure 2. Gate switch test. A) A sample gate change between XOR and NOR. B) A sample gate change between XNOR and NAND. C) A systematic test for all possible gate changes ( row: original gates, column: switched gate). All "O"'s represent a valid gate. X represents a gate that needs a global reset. All gates use the parameters from Supplementary Table 7. Given the computational complexity, we did not do an exhaustive sweep.

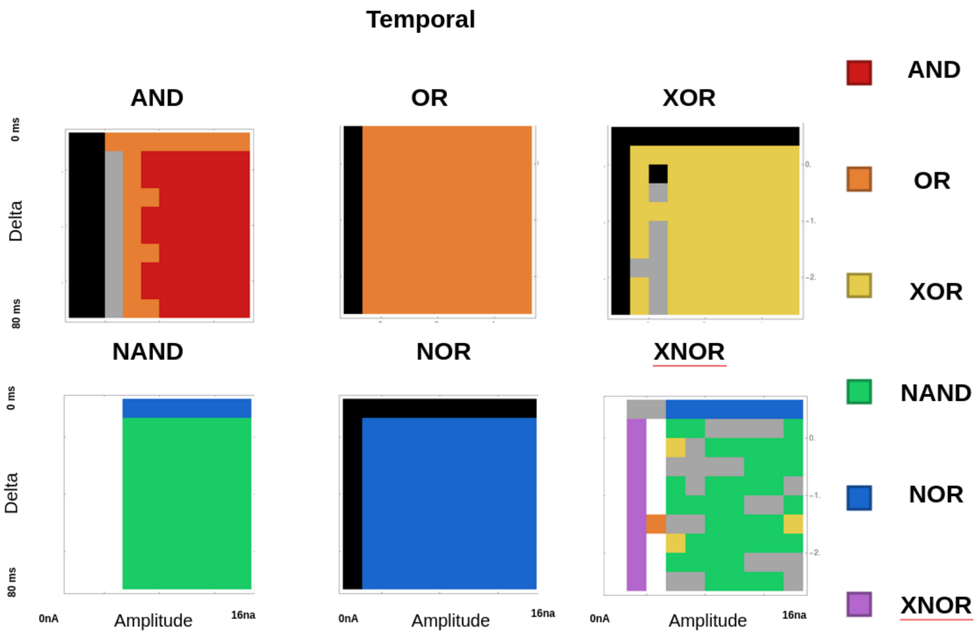


Supplementary Figure 3. A graph of robustness for the 6 symmetric temporal logic gates as a function of amplitude and the timing difference. Most gates are very robust, except for XNOR.

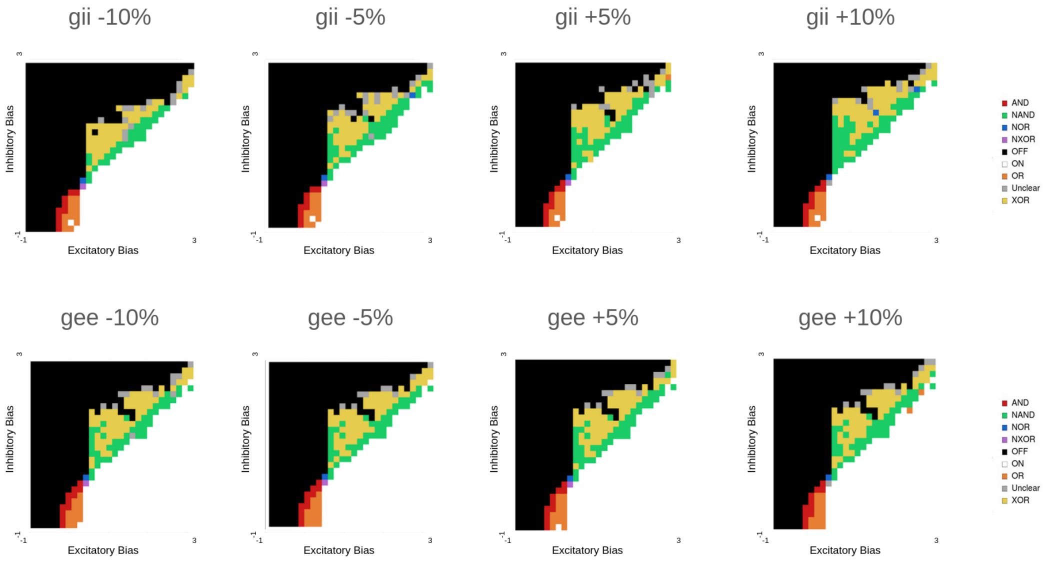


Supplementary Figure 4. Modification of synaptic weights inhibitory to inhibitory (inh) and excitatory to excitatory gee with up to a 10% perturbation both up and down. All gates exist for −10% synaptic weight perturbation. 5 out of 6 gates exist for +10% synaptic weight perturbation. XNOR is the only exception and exists up to 5% pertubration.

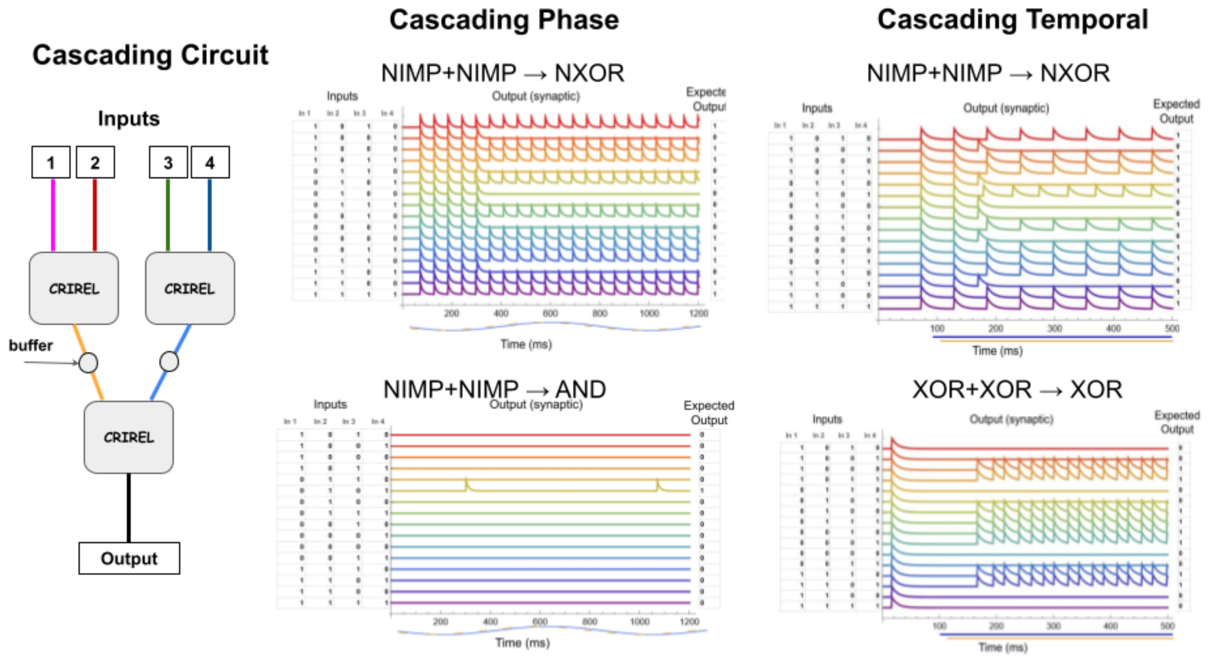


Supplementary Figure 5. The cascading circuit example for both phase and temporal computation (using Izhikevich neurons) involves an experimental setup with two layers of CRIREL circuits. Layer 1 consists of two CRIREL circuits, each taking two inputs. These inputs are either all phase inputs or temporal inputs, as defined earlier. Each CRIREL in Layer 1 outputs to a buffer neuron, which is responsible for mapping the output neuron's spikes to a fixed frequency and synaptic strength (refer to the implementation of the adder neuron). Layer 2 contains another CRIREL circuit. This CRIREL takes the previous two outputs as inputs and performs a different logical operation on them than the circuits in Layer 1, returning a new output. In this way, logic can cascade from one neuron to the next. As examples of cascading logic, we demonstrate that it can handle magnitude-type logic (full adder), phase logic (Top: NIMP + NIMP → NXOR, Bottom: NIMP + NIMP → AND), and temporal logic (Top: NIMP + NIMP → NXOR, Bottom: XOR + XOR → XOR). The parameters for these examples are the same as in previous Izhikevich neuron examples. The buffer neuron copies the parameters of the output neuron. The synaptic weight from the output neuron to the buffer neuron is set to 4.0, while the synaptic weight from the buffer neuron to the 2-layer CRIREL is set to 1.0.
